# Cell Specific Reactivation of Epicardium at the Origin of Fibro-Fatty Remodeling of the Atrial Myocardium

**DOI:** 10.1101/589705

**Authors:** Nadine Suffee, Thomas Moore-Morris, Nathalie Mougenot, Gilles Dilanian, Myriam Berthet, Bernd Jagla, Julie Proukhnitzky, Pascal Le Prince, David A Tregouet, Michel Pucéat, Stéphane N Hatem

**Author notes:** MP and SNH joined senior authors. Correspondance: Stephane Hatem, UMR_S1166, Faculté de médecine, 91, boulevard de l’hôpital, 75013 Paris, France.

## Abstract

Epicardium, the mesothelium covering the heart, is composed of multipotent cells and is reactivated following myocardial injury in adults. Herein, we provide evidence for activation of atrial epicardium in aged patients with diseased atria and in murine models of atrial remodeling. Epicardial activation contributed to fibro-fatty infiltration of sub-epicardium that contained a number of cells co-expressing markers of epicardial progenitors and fibroblasts. Indeed, using genetic lineage tracing of adult epicardium, we demonstrate the epicardial origin of fibroblasts within fibro-fatty infiltrates. A subpopulation of adult epicardial-derived cells (aEPDCs) expressing PDGFRα, niched in the sub-epicardium, were isolated and differentiated into myofibroblast in the presence of angiotensin-II. Furthermore, single cell RNA-seq analysis identified several clusters of aEPDCs and revealed transition from adipogenic to fibrogenic cells. In conclusion, a subset of aEPDCs, pre-programmed towards a specific cell fate, contributes to fibro-fatty infiltration of sub-epicardium of diseased atria.

## Introduction

The epicardium plays a major role in the formation of embryonic heart. This outer mesothelial layer of the heart contains multipotent progenitors, giving rise to smooth muscle cells of coronary vessels as well as to cardiac fibroblasts that provide a stroma that drives formation and alignment of myocardial fibers^1^. The epicardium is quiescent in healthy adult heart, but can be reactivated following massive and acute injury. It thickens, expresses fetal genes, and triggers differentiation of epicardial progenitors that can migrate into subjacent myocardium. Hence, it has been shown that the epicardium can be a source of fibroblasts contributing to fibrosis of the adult ventricular myocardium in response to acute ischemia^2^. In addition, we and Yamaguchi *et al* have reported that atrial epicardial cells undergo epithelial-to-mesenchymal transition (EMT), delaminate from the epicardium and differentiate into adipocytes contributing to the accumulation of epicardial adipose tissue (EAT) both at the external face of the epicardium and in the sub-epicardium ^3, 4^.

In the atria, EAT is a common component of atrial histology. It is a source of free fatty acid and of adipokines that can regulate the metabolism of the redox state of neighboring myocardium^5, 6^. However, both expansion and fibrotic remodeling of EAT are associated with an increased risk of atrial fibrillation (AF), the most frequent cardiac arrhythmia in clinical practice^7^. One mechanism linking fibro-fatty infiltration of the subepicardium and AF is that this remodeling process contributes to the electrical dissociation between subepicardium and subendocardium areas favoring focal fibrillation waves as recorded in patients with longstanding AF^8, 9^. Hence, fibro fatty infiltration of the subepicardium is considered as an important determinant of AF substrate. However, the underlying biological mechanisms are still largely unknown.

The natriuretic peptide secreted by stretched atrial myocytes is a potent adipogenic factor for adult epicardium progenitor-derived cells (aEPDCs) which might favor EAT expansion in the context of increased atrial workload^4^. EAT can contribute to the fibrotic remodeling of the subepicardium by secreting adipo-fibrokines such as activin-A that freely diffuse into the neighboring myocardium^10^. In addition, the adipose tissue depot commonly seen in the subepicardium can be replaced by fibrosis through an immune process mediated by CD8+ T lymphocytes^11^. Altogether these results suggest a balance between EAT expansion and fibrosis of the subepicardium.

In the present study, we tested the hypothesis that the atrial epicardium can be a source of fibroblasts contributing to the fibro-fatty infiltration of the subepicardium. We found that a subset of aEPDCs are programmed towards fibroblast or adipocyte differentiation, and contribute to fibro-fatty infiltration of the subepicardium in response to various stimuli. Our results provide novel insight into the role played by the epicardium in the slow process of atrial remodeling and the formation of AF substrate.

## Methods

### Study approval

All animal experiments were conformed to the Guide for the care and use of laboratory animals, according to the Directive 2010/63/EU of the European Parliament approved by the local committee of animal care (agreement A751315).

Human tissue samples were obtained from patients that had undergone heart surgery. Data and samples were obtained in accordance with French Law “loi Huriet-Sérusclat” and with the approval of the Ethical Committee (Comité de protection du personnel Ile-de-France VI) of Pitié-Salpétrière Hospital non-opposition to the research was obtained from each patient. Personal Data treatment necessary to the research was declared to the National Commission for Data Protection and Liberties (CNIL-France) under the Data Protection Act number 78-17. Clinical parameters are indicated in Supplementary Table 1.

### Murine model

Both, eight-week-old male mice and rats were maintained under 12 h light/ 12 h dark cycle at constant temperature (21°C) with free access to food and water. Embryos were staged as E11.5 or E16.5 days. WT1Cre*^ERT2+/-^* and Rosa26*^tdTomato+/+^* on a C57BL/6J background were purchased from The Jackson laboratories (L’arbresle, France). WT1Cre*^ERT2+/-^*mice were bred with Rosa26*^tdT+/+^* mice as described previously^4^. In order to label adult epicardium, 6 week-old WT1Cre*^ERT2+/-^*;Rosa26*^tdT+/-^* mice were administered tamoxifen (70mg/kg bw) for 5 consecutive days. Wistar rats were purchased from Janvier (200-220 g, CERJ, Laval, France). WT1Cre*^ERT2+/-^*;Rosa26*^tdT+/-^* transgenic mice (n=10) and Wistar rats (n=50) were anesthetized with Isoflurane 2 %. Mice and rats underwent thoracotomy and definitive occlusion of the left anterior descending coronary artery was either left intact (sham) or occluded with a 5.0 nylon monofilament suture at 1 mm from its origin to cause myocardial infarction (MI).

### Clinical study of experimental models

Cardiomyopathy development in rat and mouse was monitored under 1.5-2 % isoflurane anesthesia by transthoracic echocardiography as previously described^12^, using a Vivid 7 dimension cardiovascular ultrasound system equipped with a probe of 9-14 MHz frequency (GE Healthcare, Vélizy, France). Bi-dimensional (B) and time-motion (TM) mode views were recorded along the parasternal long axis from mitral and aortic valves to apex and in a short axis at the level of the mid papillary muscles. Left atrial area (LAA) was measured using B-mode. TM-mode was used for measurements of left atrial diameter and left ventricle parameters such as diameter (Vd) and volume (Vd) at end diastole, diameter (Ds) and volume (Vs) at end systole. The percentage of left ventricular fractional shortening (FS) in the long-axis view was calculated as (Dd – Ds)/Dd x 100. The left ventricle ejection fraction percentage (EF) was calculated as (Vd-Vs)/Vd x 100. All measurements were performed and averaged over five consecutive cardiac cycles by one single blinded operator. Reproducibility was assessed by triplicate measurements.

Atrial fibrillation episodes were induced by atrial burst pacing. Under anesthesia, a 4-French stimulation catheter (Saint Jude medical) was inserted into the esophagus down to the level of the left atria. Left atrial was then stimulated for 3 seconds at 10 Hz and an electrocardiogram (ECG) was recorded. AF duration was calculated as the time between the end of stimulation and recovery of sinus rhythm.

### Human tissue

Appendage samples of human atrial tissue were obtained for secretome studies (n=10), fixed and embedded in paraffin (n=109) for histology and immunofluorescence analysis or were used for the isolation of epicardial progenitor cells (n=15) as previously described (Supplementary Table 1)^4^.

### Histology

Mouse and rat hearts were removed, perfused through the aorta with PBS, fixed in 4 % paraformaldehyde (PFA) overnight. Mouse hearts were then dehydrated overnight in sucrose and embedded in OCT and frozen as described previously^4^. Human atrial appendage samples or rat hearts were fixed, dehydrated and paraffin embedded. Seven-µm-thick sections atria samples were stained with Masson’s trichrome according to manufacturer’s instructions (Sigma-aldrich). Bright field images were acquired with a Nikon DS-Ri1 camera coupled to Eclipse-Ti Nikon microscope and Nis-Element software (Nikon France S.A.) and analyzed with Image J software.

### Histomorphometry

Seven-µm-thick sections of human or murine atrial samples were stained with PicroSirius red according to manufacturer’s instructions (Sigma-aldrich). Bright images were acquired with a Nikon DS-Ri1 camera coupled to Eclipse-Ti Nikon microscope and Nis-Element software (Nikon France S.A.) and analyzed with Histolab software (Evry, France). Collagen type I and III were visualized with polarization microscopy. The identification of fibers was based on the birefringence of collagen that was modulated by the binding of PicroSirius molecules. Polarization images were acquired with a Zeiss AxioObserver Z1 (Carl Zeiss, Germany) microscope, using a 20x NA 0.8 oil immersion objective and a Zeiss HRc camera. For image acquisition we used ZEN Blue software (Carl Zeiss, Germany).

### Differentiation of aEPDCs

Human aEPDCs were culture in basal DMEM/M199 (1:1) and were differentiated into adipocytes with ANP (10 pM) for 21 days as described previously^4^. To induce human aEPDC-derived fibroblasts, progenitors were incubated in basal medium supplemented with Ang-II (10 nM) or TGF-β (10 nM) (Sigma-Aldrich) for 7 or 21 days. A fibrogenic medium was used as a positive control, composed of DMEM/F12 (Invitrogen) supplemented with L-glutamin (2 mM), FCS (10 %) (Sigma-Aldrich), b-FGF (10 ng/ml), TGF-β (10 nM) and penicillin-streptomycin (1%) (ThermoFisher Scientific)^13–15^. All media were changed twice per week.

### Immunofluorescence assay

Adult EPDC-derived fibroblasts, human or murine atria tissue sections were incubated overnight with antibodies against collagen-1 (ab34710), αSMA (ab7817), PDGFRα (ab5460), DDR2 (ab126773), FSP-1 (ab27957), WT1 (ab96792), Pref1 (ab119930), fibronectin (ab2413), Perilipin (ab3526), AGTR1 (ab124505), NPRA (ab70848) or Tcf21 (ab32981) purchased from Abcam and vimentin (#5741) or CD44 (#3570) purchased from Ozyme. Then antibodies were revealed using Alexa-Fluor 488 or Alexa-Fluor 594 secondary antibodies (Thermo Fisher Scientific). Sections also were incubated with DAPI solution (Sigma-Aldrich). Images were acquired with a Nikon DS-Ri1 camera coupled to an Eclipse-Ti Nikon microscope and Nis-Element software (Nikon France S.A.) or Zeiss LSM800 (Carl Zeiss, Germany) and analyzed with Image J software.

### Oil red O staining

Adult EPDC-derived cells were fixed with PFA (4 %) and stained with oil-red-O as described previously^4^. All images were acquired with a Nikon DS-Ri1 camera coupled to an Eclipse-Ti Nikon microscope and Nis-Element software (Nikon France S.A.), and analyzed with Image J software.

### Flow Cytometry and cell sorting

Characterization of human aEPDCs was assessed by using 1.10^6^ cells/mL. Cells were passaged using TrypLE (Applied Biosystems) and were washed with PBS containing 5 % (vol/vol) FCS as described previously.

Human aEPDCs were incubated with specific membrane antibodies against PDGFRα (ab5460), DDR2 (ab126773), Pref1 (ab21682; Abcam), Perilipin (ab3526; Abcam), AGTR1 (ab124505) or NPRA (ab70848) purchased from Abcam, and their specific isotype controls for 30 min at room temperature. Cells then were incubated with Alexa-Fluor 650, washed, fixed with 0.1 % PFA, and analyzed by flow cytometry using a MACSQuant analyzer (Miltenyi Biotech, Paris, France), and FlowJo software (FlowJo LLC). Cells sorting was performed with coupled antibodies against PDGFRα-FITC or Pref1-APC. Cells were sorted by ASTRIOS (Beckman Coulter, Villepinte, France) and were cultured in aEPDC culture medium.

### Single cell RNA-sequencing

Adult EPDCs from sham or HF rats were isolated from atrial epicardium explants and passaged once in culture. Epicardial cells and aEPDC in culture were dissociated with trypsin into a single cell suspension in the presence of 100 nM thiazovivin, filtered through a 40 μ m mesh and 10 000 cells were processed with the SingleCell3 Reagent Kit on the Chromium platform as described by the manufacturer (10X genomics). cDNA libraries were sequenced with a Next-seq Illumina sequencer. A first analysis was performed with cell Ranger and C-loop 10X genomics softwares. Clusters and subclusters were defined using genes differentially expressed in cell clusters in comparison with all other cells with a threshold of Log2 equal to at least 2. Then a secondary analysis was done using the computional workflow ScshinyHub (https://github.com/baj12/scShinyHub), developed by B. Jagla at Pasteur Institute, Paris. A total number of 16 400 genes were detected and an average of 3400 genes was expressed in all cells. Trajectory inference was performed using SCORPIUS package within ScshinyHub.

### RNA Extraction and Reverse Transcription and qPCR Assay

Total RNA was isolated from 1.10^4^ aEPDCs-derived fibroblasts and qPCR was performed using a TaqMan Gene Expression Assay as described previously^4^ (Supplementary Table 2). Expression relative to GAPDH was calculated using the 2-(ΔΔCt) method. Gene expression levels were expressed in arbitrary units (AU) and were normalized to control.

### Western Blot Analysis

Human aEPDCs were cultured in six-well plates and were treated with fibrogenic medium or Ang-II (10 nM) for 7 days. Cellular lysis was performed as described previously^4^. Proteins (10 μg) were separated on 10 % (vol/vol) SDS-polyacrylamide gels followed by electrophoretic transfer of protein to nitrocellulose membrane (Thermo Fisher Scientific). Membranes were incubated overnight with antibodies against Smad2/3 (#8685 and #3122) or p38-MAPK (#4511 and #9212) (Ozyme). Expression levels relative to GAPDH (# 2118; Ozyme) were expressed in arbitrary units and were normalized to control. Membranes were revealed using streptavidin-HRP–conjugated secondary antibody solution (Thermo Fisher Scientific). Membranes were exposed to an Image Quant LAS4000 camera (GE Healthcare) and were analyzed with Image J software.

### Statistics

Data obtained are expressed as means ± S.E.M. Differences were investigated using un-paired 2-tailed Student’s *t*-test or a one-way analysis of variance (ANOVA) with Bonfferoni’s post-hoc test, considered significant at *P* < 0.05. Statistical analysis was performed with Prism GraphPad 6.0 (Software Inc.).

Before any statistical analyses of histomorphometry, log-transformation was applied to reduce any skewness in the distribution of the epicardial adipose tissue and epicardial fibrosis variables. Pearson correlation coefficients were computed between biological and anthropometric variables while adjusting for age, gender and BMI when appropriate. Association of biological parameters with clinical outcomes was assessed using logistic regression adjusted for age and gender. The statistical threshold used for declaring statistical association was fixed to 0.05. Analyses were performed using the R statistical computing software.

## Results

### Evidence for epicardium activation and remodeling in human atria

In human right atrial tissue sections, the epicardium presented various histological aspects. These included a thin cell monolayer separated from adjacent myocardium (Fig. 1a, (i)) or adipose tissue (Fig. 1a, (ii)) by a regularly arranged basal extracellular matrix (ECM) mainly composed of collagen-1 (Fig. 1a, (iii)). Alternatively, it constituted a thick layer (47.43 ± 17.7 mm²) in continuity with dense fibrosis (Fig. 1a, (iii,iv)). Under polarized light, extracellular matrix of the thin epicardium was mainly composed of regularly arranged yellow-red strong birefringence collagen-1 (Fig. 1b, (ii)), whereas in thick and fibrotic epicardium, collagen-1 and −3 (yellow-green birefringence) fibers were disorganized (Fig. 1b, (iii,iv))^16^. A number of cells co-expressing mesenchymal cell markers αSMA (8.4 ± 0.9%) or PDGFRα (10.3 ± 2.1 %) were present in the thick and fibrotic epicardium (Fig. 1c, Supplementary Table 3a).

**Fig. 1.**
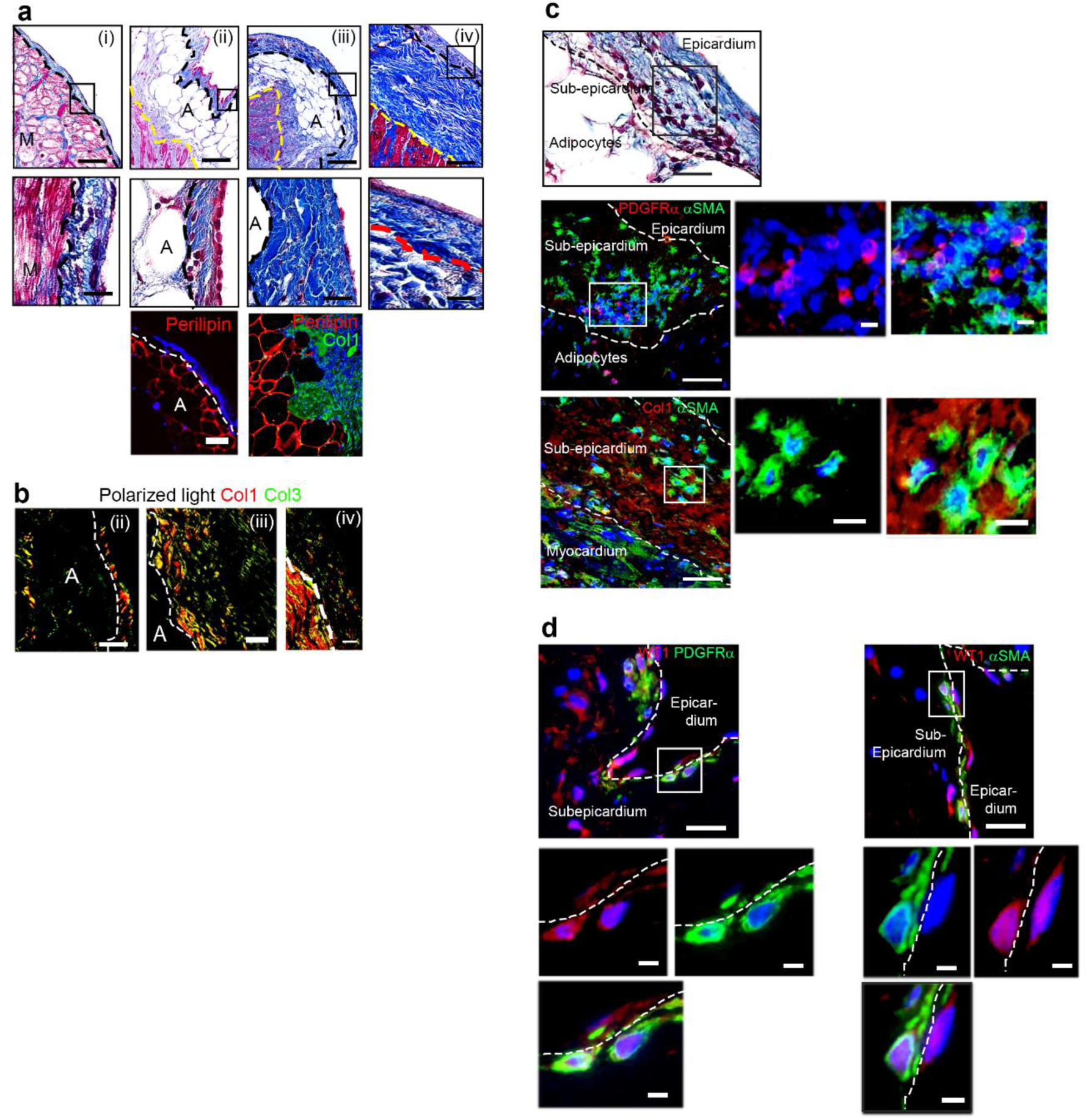
Epicardium progenitor profile in human atrial epicardium expansion. (**a**,**b**) Seven-µm-thick sections of human atrial tissue (n=109) stained with Masson’s trichrome and observed by transmitted light or (**a**) or PicroSirius red observed with polarized light microscopy (**b**) (collagen-1, Col1, red-yellow; collagen-3, Col3, green). Scale bar, 50 µm, 25 µm, inset 10 µm. (**b**-**d**) Immunofluorescence staining of human atrial tissue sections (n=6) for Perilipin-1 (**a**), Col1 (**a**,**c**), βSMA (**c**,**d**), PDGFRβ (**c**,**d**) and WT1 (**d**). Scale bar, 100 µm, 25 µm, 10 µm. M, myocytes; A, adipocytes.

The percentage of fatty infiltrates correlated negatively with the epicardium thickness (−0.52, P<0.01, n=109) suggesting a mutual exclusion of the two histological components. Hence, the ratio of adipose tissue to epicardial fibrosis (EAT/epi Fibrosis) was used as a marker of epicardial area remodeling. Looking deeply into these data revealed that EAT/epi Fibrosis tended to differentially associate with clinical outcomes according to patient age. Indeed, in patients over 70 years-old (which corresponds to the median age in our population), we observed that the EAT/epi Fibrosis ratio significantly decreased in atrial sections from patients with valve mitral diseases (0.33 vs 0.84, p < 0.05) and that there was a tendency to decrease in patients with permanent AF (0.32 vs 0.86, p = 0.211) with no overlap between both groups (Supplementary Table 4). These histological observations suggest that low grade activation of the epicardium can contribute to the remodeling of the atrial myocardium.

### Recruitment of epicardial progenitors into the subepicardium of human atria

We previously reported the presence of cells expressing the marker of epicardial progenitors Wilm’s tumor (WT)-1 and of pre-adipocytes in the subepicardium of human atria^4^. Here, we found that WT-1 positive cells expressing markers of myofibroblasts (αSMA) or fibroblasts (PDGFRα) too were detected in the subepicardium of human atria (Fig. 1d) (Supplementary Table 3a).

These data raised the question as to the presence of subset of cells derived from epicardial progenitors and that could differentiate into distinct mesenchymal cell lines *i.e* adipocyte or fibroblasts. This idea was further supported by the observation that atrial aEPDCs showed distinct behavior in culture when treated with a myocardial secretome from human atria. After 21 days in culture, one-third of the cells presented fibroblast phenotypes including an elongated shape and expression of ECM proteins such as collagen-1 and fibronectin together with fibroblast markers such as vimentin, αSMA, PDGFRα and DDR2. The remaining two-thirds showed an adipocyte phenotype, expressing perilipin and containing lipid droplets (Fig. 2a-c).

**Fig. 2.**
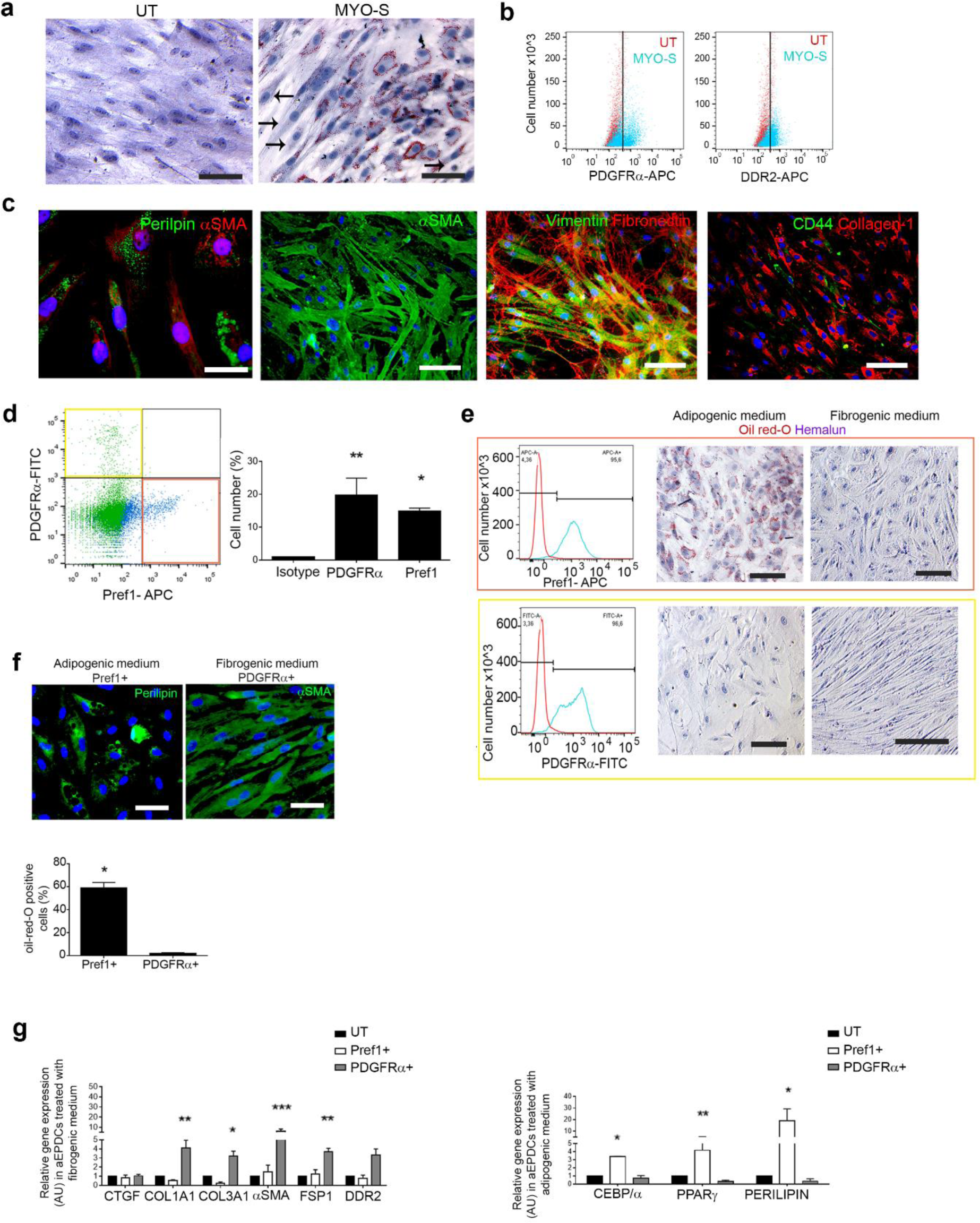
Fibrogenic or adipogenic fates of aEPDCs. (**a**) Representative images of aEPDCs incubated with MYO-S stained in oil-red-O and counterstained with hematoxylin (n=10). Scale bar, 100 µm. Black arrows indicate elongated shape cells. (**b**) Flow cytometry analysis of aEPDCs treated with MYO-S (n=10). (**c**) Immunofluorescence staining in aEPDCs treated with MYO-S for Perilipin, αSMA, Vimentin, Fibronectin, CD44 and Collagen-1. Scale bars, 100 µm, 50 µm. (**d**) Analysis of PDGFRα and Pref1 expression in aEPDCs by flow cytometry (n=10). (**e**) Representative images and histogram of sorted PDGFRα^+^ or Pref1^+^ aEPDCa incubated with adipogenic medium or fibrogenic medium and stained with oil-red-O and counterstained with hemalun (n=10). Scale bars, 100 µm, 50 µm. (**f**) Perilipin or αSMA immunofluorescence staining of PDGFRα^+^ or Pref1^+^ sorted aEPDCs cultured with adipogenic medium or fibrogenic medium. Scale bar, 50µm. (**g**) Transcript expression levels of adipogenic and fibrogenic markers in PDGFRα^+^ or Pref1^+^ aEPDCs treated with fibrogenesis (left) or adipogenesis (right) medium.

Next, aEPDCs were sorted based on the expression of Pref1 or PDGFRα (Fig. 2d). After 21 days of culture in the presence of a fibrogenic medium, only PDGFRα^+^ cells presented fibroblast phenotypes with αSMA labeling (Fig. 2e,f). In contrast, in the presence of an adipogenic medium, only Pref1^+^ cells became round-shaped and contained oil red-O stained lipid droplets and expressed perilipin typical of adipocytes (Fig. 2e,f). In addition, fibrogenic gene expression was observed in PDGFRα^+^ cells when cultured in fibrogenic medium, whereas Pref1^+^ cells cultured under adipogenic medium induced expression of adipocyte transcripts (Fig. 2g). These results suggest that distinct populations of aEPDCs can differentiate into fibroblast or adipocytes in response to various hormonal stimulations.

### Distinct signaling pathways regulate differentiation of subsets of human aEPDCs

Next, we attempted to identify if distinct signaling pathways could regulate aEPDC fate. Following our previous finding of the presence of the adipogenic factor ANP in myocardial secretome (MYO-S)^4^, we hypothesized that fibrogenic factors were also present in the secretome. Several candidates were detected in MYO-S such as TGF-β or Angiotensin (Ang)-II^4^. Interestingly, *in vitro*, aEPDCs acquired a fibroblast phenotype in the presence of Ang-II or TGF-β, as indicated by the up-regulation of PDGFRα and DDR2 (Fig. 3a, Supplementary Fig. 1a). In addition, at the transcript level, expression of the various ECM and fibroblast genes were up-regulated (*COL1A1*, *CTGF*, *TGFB*, *PDGFR*α and *FSP1)* whereas that of adipogenic (*CEBP/*α, *PPARγ*, *PERILIPIN-1*) genes were down-regulated in Ang-II treated cells compared with ANP treated cells (Fig. 3b,c, Supplementary Fig. 1b). Moreover, Ang-II inhibited perilipin-1 protein expression compared to ANP (Fig. 3d) and a subset of aEPDCs remained Pref-1^+^ (Fig. 3e). Finally, the fibrogenic canonic Smad2/3 and p38-MAPK signaling pathways were activated in this culture condition (Fig. 3f)^17, 18^.

**Fig. 3.**
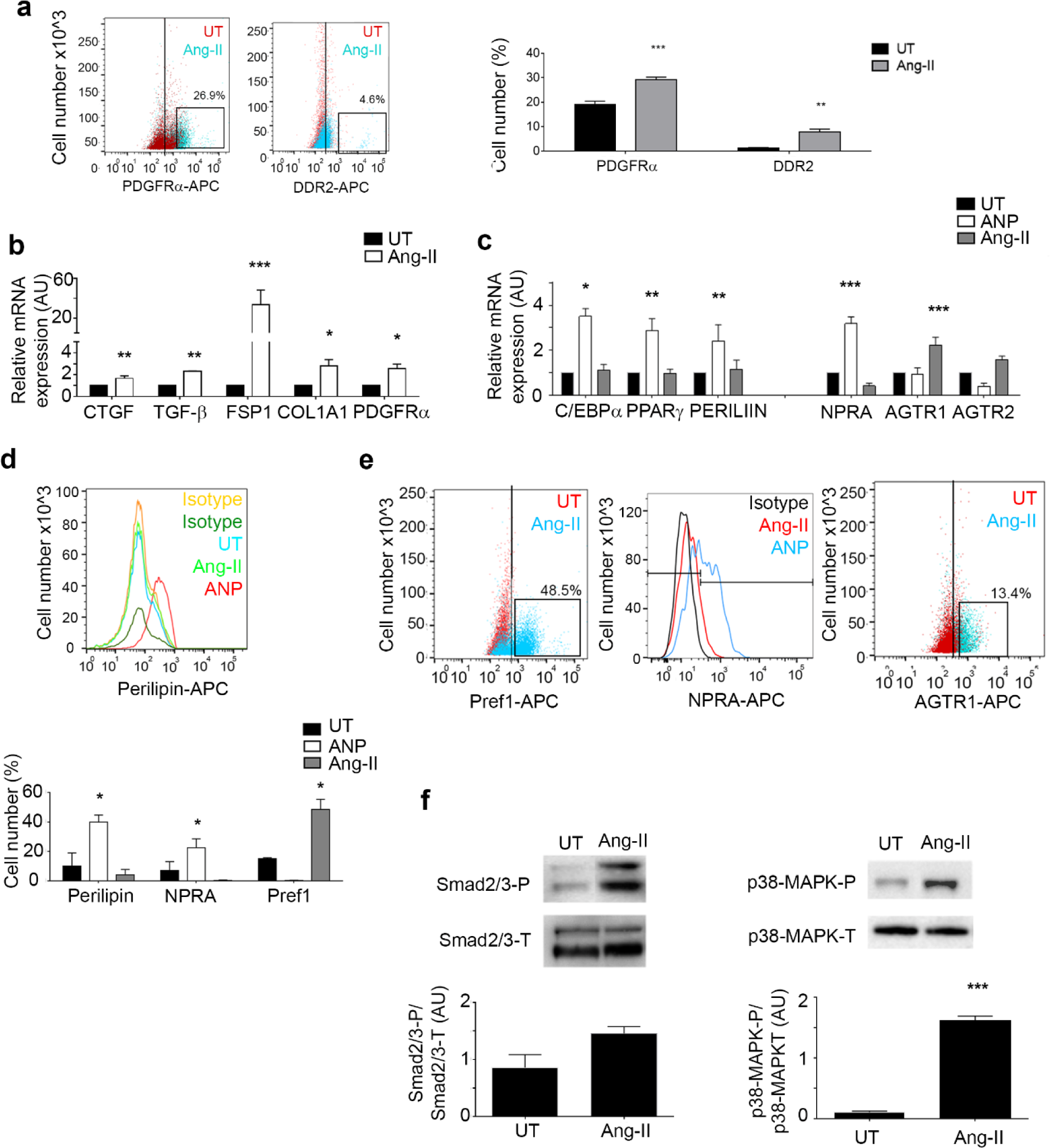
Angiotensin-II induces fibrogenic differentiation of aEPDCs. **(a)** Flow cytometry and histogram analysis of aEPDCs treated with Ang-II (10 nM) for 7 days. **(b,c)** Transcript expression levels of fibrogenic **(b)**, adipogenic, ANP or Angiotensin-II receptors markers **(c)** in aEPDC treated with Ang-II (10 nM) **(b,c)** or ANP (10 pM) for 21 days **(c)**. **(d,e)** Flow cytometry analysis and quantification of aEPDCs treated with Ang-II (10 nM) or ANP (10 pM). **(f)** Immunoblots and relative quantification of Smad2/3 or p38-MAPK signaling pathways in aEPDCs treated with Ang-II (10 nM) for 21 days. Data are expressed as mean ± SEM of n=4 independent experiments. *, P < 0.05, **, P <0.01, *** P <0.001, one-way ANOVA with Bonferroni’s post hoc test.

Interestingly, both transcript levels and the plasma membrane expression of Angiotensin-II receptor 1 (AGTR1) were induced by Ang-II in aEPDCs, whereas the ANP receptor, NPRA, was undetectable (Fig. 3d,e). The opposite result was obtained in cells treated with ANP *i.e* up-regulation of NPRA and down regulation of AGTR1 expression were observed (Fig. 3d,e). Flow cytometry showed that Ang-II increased the fraction of cells expressing PDGFRα and the pre-adipocyte marker, Pref1, whereas cell expressing perilipin, *i.e.* mature adipocytes, could not be detected (Fig. 3a, 3e). These results indicate that distinct signaling pathways regulate aEPDC differentiation.

### Adult EPDC are recruited of at early stage of experimental atrial remodeling

To examine whether the epicardium is activated during the atrial remodeling, we studied a well-characterized model of atrial remodeling secondary to ischemic HF in rats^19^. Two months following the onset of the ischemic cardiopathy, all animals were in HF with dilated atria (Fig. 4a,b, and Supplementary Fig. 2a-c). They showed a vulnerability to AF as indicated by a longer duration of triggered episodes of AF compared to shams (Supplementary Fig. 2d,e). After sacrifice, histological analysis revealed a hypertrophied and fibrotic myocardium in dilated atria of HF rats compared to sham (epicardial fibrosis area in atria 70.3 ± 8.3 % vs 42.8.2 ± 7.9 %, P < 0.001) (Fig. 4a,b). In addition, collagen-1 and collagen-3 were also disorganized and irregularly arranged (Fig. 4c).

**Fig. 4.**
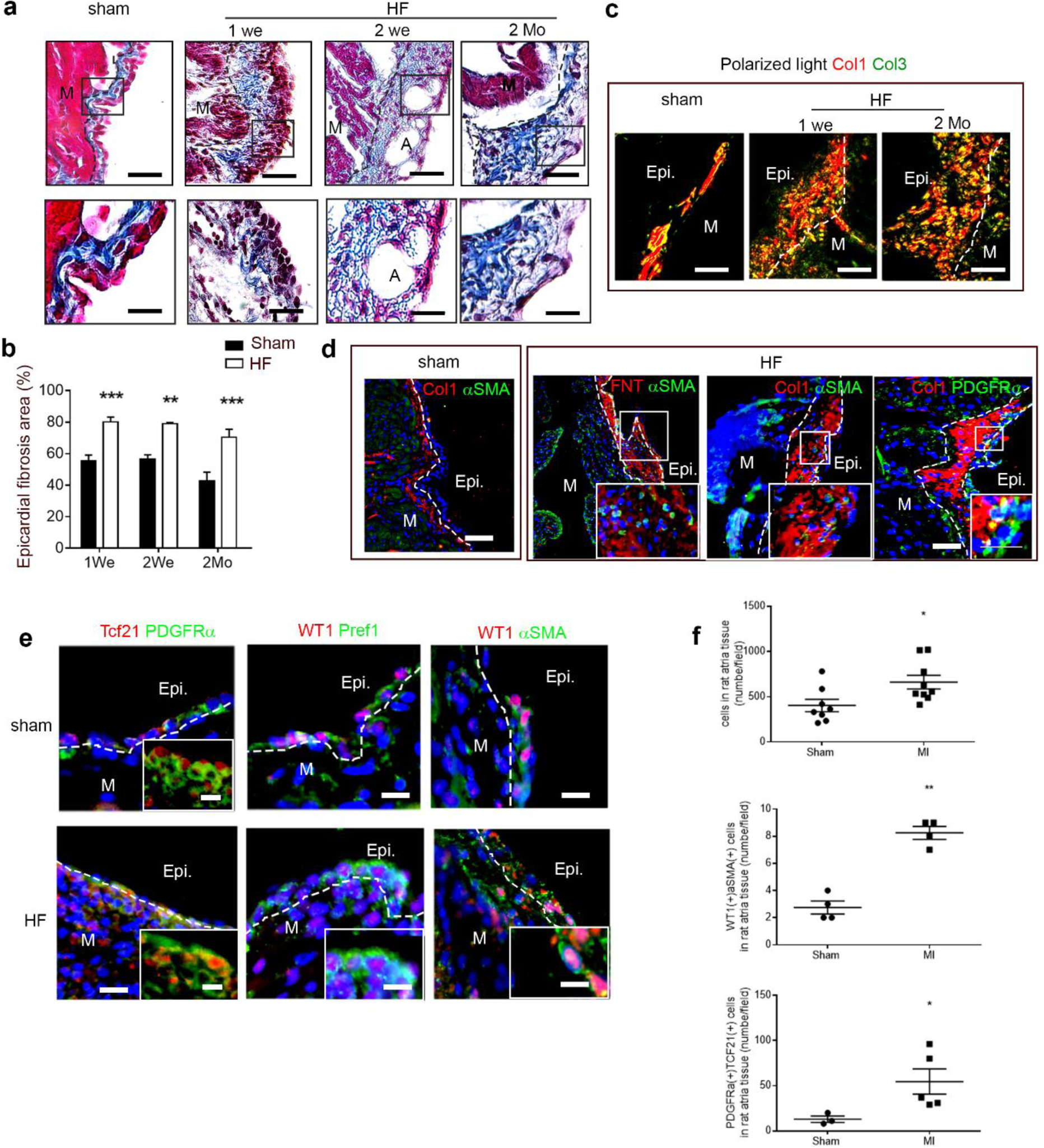
Epicardium remodeling is an early event during atrial remodeling in rat model of heart failure. (**a**) Masson’s trichrome staining of 7-µm-thick sections of rat atrial tissue at 1 week, 2 weeks or 2 months post-MI and Sham (n=5, each group) observed by transmitted light microscopy. Scale bar, 50 µm, 25 µm. (**b**) Histomorphometry assay quantification in rat atrial tissue post-MI (n=5, each group) and Sham (n=5, each group). (**c**) PicroSirius red of 7-µm-thick sections of rat atrial tissue at 1week or 2 months post-MI and Sham (n=3, each group) is observed under polarized light microscopy, collagen-1 (Col1, red-yellow), collagen-3 (Col3, green). Scale bar, 60 µm. (**d**,**e**) Immunofluorescence staining in rat atrial tissue sections at 2 months post-MI (**d**), 1 week post-MI (**e**) or Sham for Col1, αSMA, PDGFRα (**d**,**e**), fibronectin (FNT) (**d**), WT1, Pref1 and Tcf21 (**e**). Scale bars, 200 µm, 100 µm, 50µm. (**f**) Scatter plots represent number of cell per field in sub- or epicardium of rat atria tissue. Data are expressed as mean ± SEM of at least n=4 independent experiments. *, P < 0.05, **, P < 0.01, *** P <0.001, one-way ANOVA and Bonferroni’s post hoc test. Epi., epicardium; A, adipocytes; M, Myocytes.

Interestingly, the epicardium was thick (epicardial fibrosis area in atria 78.95 ± 1.6 % vs 56.61 ± 4%, p < 0.001) and a packed fibrosis composed of collagen and fibronectin depots was observed in the subepicardium of dilated atria of HF rat (Fig. 4d). Both fibroblasts expressing PDGFRα and myofibroblasts expressing αSMA were detected in the subepicardial fibrosis (Fig. 4d,e). In addition, adipocytes were observed (epicardial adipocyte area in atria 37.72 ± 6.13 %, P < 0.05) (Fig. 4a). Expansion of the epicardium was observed already one week after MI (80.12 ± 8.6 % vs 55.25 ± 7.1 %, p < 0.01), and reached a plateau with progression of the cardiomyopathy (Fig. 4a,b). Some cells were double-positive for αSMA and WT1 or PDGFRα and Tcf21, a transcription factor involved in fibroblast cell fate acquisition^20^ (Fig. 4e,f, Supplementary Table 3b). Interestingly, some WT1(+); Pref-1(+) cells were localized in epicardial monolayer (Fig. 4e,f, Supplementary Table 3b). These results indicate that epicardium expansion is an early event in the development of atrial cardiomyopathy and that fibroblasts derived from activated epicardium could contribute to the remodeling of the atrial subepicardium.

### Evidence for epicardial progenitor origin of fibroblasts in the atrial subepicardium

In order to obtain direct evidence that epicardium could be a source of fibroblasts during atrial remodeling, we reproduced the model of atrial cardiomyopathy secondary to ischemic HF in the WT1Cre*^ERT2+/-^*;ROSA26*^tdT+/-^* mice, allowing genetic lineage tracing of WT1-positive epicardial progenitor cells in adult heart^2, 4, 21, 22^. As in rats, two months after surgical MI, mice had developed HF with dilated and remodeled atria (Supplementary Fig. 2f-h). Furthermore, the epicardium was thick and remodeled (epicardial fibrosis area in atria 78.13 ± 5.5 vs 53.46 ± 4.5, p < 0.001) with packed and dense fibrosis (Fig. 5a,b).

**Fig. 5.**
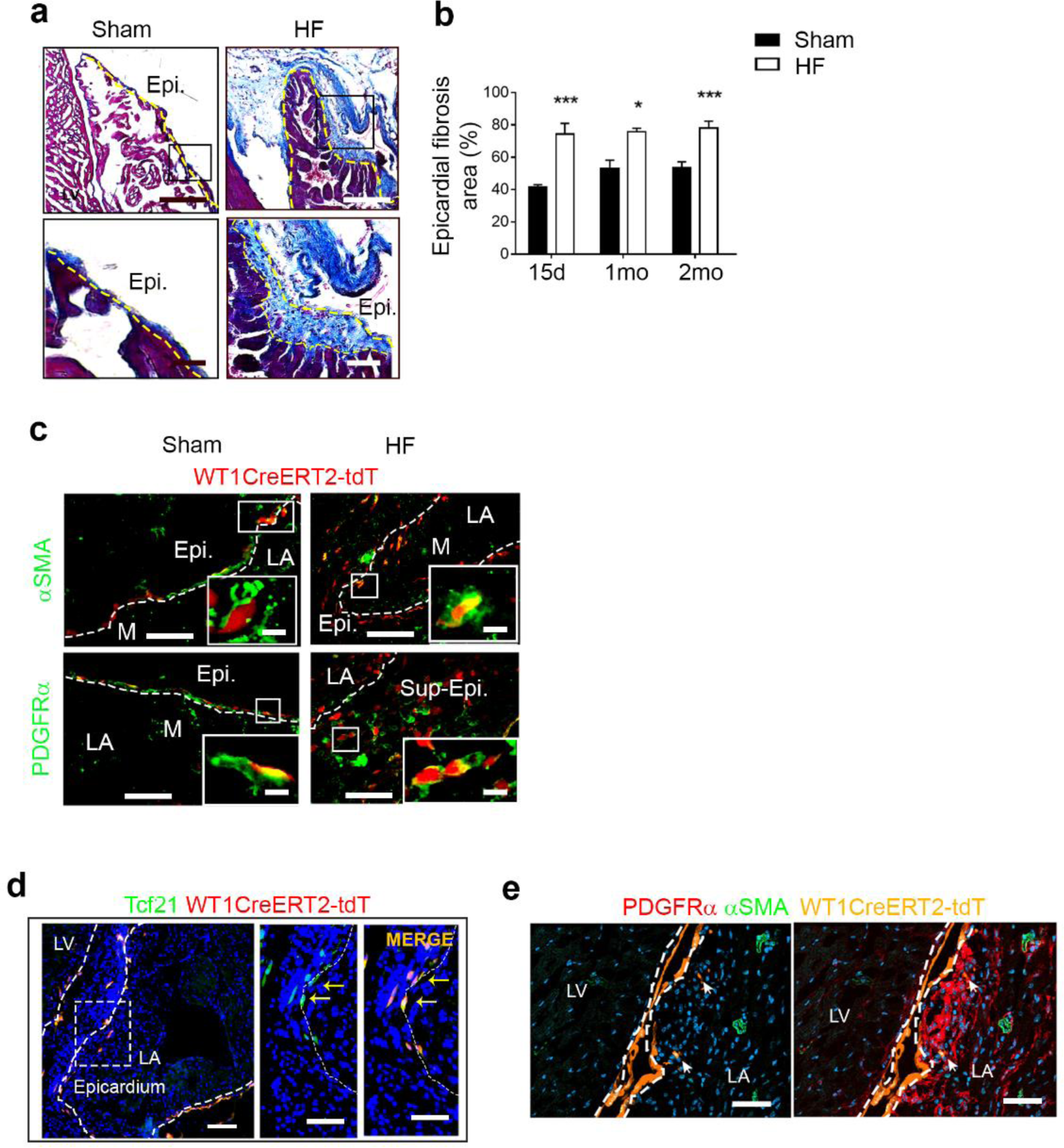
Epicardium is a source of fibroblasts during atrial remodeling in mouse model of heart failure. (**a**) Masson’s trichrome staining of 7-µm-thick sections of WT1CreERT2^+/-^/Rosa-tdT^+/-^ mouse atrial tissue at 2 months post-MI or Sham (n=5, each group). Scale bar, 200 µm, 100 µm. (**b**) Histomorphometry assay quantification in WT1CreERT2^+/-^/Rosa-tdT^+/-^ mice atrial tissue at 15 days, 1 months or 2 months post-MI (n=5) or Sham (n=5). (**c**-**e**) Immunofluorescence staining of WT1CreERT2^+/-^/Rosa-tdT^+/-^ mice atrial tissue sections at 2 months post-MI or Sham for αSMA or PDGFRα (**c**,**e**), Tcf21 (**d**). PDGFRα^+^ fibroblasts derived from adult epicardium are indicated by arrows (e). Scale bars, 200 µm, 100 µm, 50 µm. Data are expressed as the mean ± SEM of n=4 independent experiments. *, P < 0.05, *** P <0.001, one-way ANOVA and Bonferroni’s post hoc test. Epi., epicardium; Sup-Epi., sup-epicardium; LV, left ventricle; LA, left atria.

As expected tdT^+^-WT1^+^ cells predominated in the epicardial layer in both sham and HF mice (Fig. 5c). In contrast, only in the latter group, tdT^+^ cells were seen in the sup-epicardium of the atria indicating epicardial activation and aEPDC migration (Fig. 5c). Some of the WT1-positive cells of sup-epicardium expressed either αSMA or PDGFRα (Fig. 5c). Atrial epicardial progenitors in WT1Cre*^ERT2+/-^*;Rosa26*^tdT+/-^* mice also expressed the fibroblast marker Tcf21 (Fig. 5d). More limited numbers of adult epicardium derived PDGFRα^+^ fibroblasts were found in the subepicardium of the atria (Fig. 5e). Of note, epicardial-derived myofibroblasts were observed associated only with atrial epicardium but not with ventricular epicardium outside of the infarcted area (Fig. 5e). Interestingly, we observed heterogeneous expression of PDGFRα and Pref1 in embryonic and adult atrial epicardium, suggesting that epicardial cell heterogeneity may underlie aEPDC composite (Supplementary Fig. 3a). In embryo at E11.5, both Pref-1(+) and PDGFRa(+) cells were located in epicardium monolayer (Supplementary Fig. 3a). Interestingly, after EMT process (E16.5)^23^, only Pref1(+) cells were observed in epicardium whereas PDGFRa(+) were in sub-epicardium similar to sham adult atria (Supplementary Fig. 3a). In addition, in embryos, epicardium contained Pref1(+) cells was co-expressed with NPRA and PDGFRa(+) cells was co-expressed with AGTR1 (Supplementary Fig. 3b,c). These results indicate that resident atria epicardial progenitor cells reactivate to differentiate into fibroblasts or adipocyte during atrial remodeling.

### Single cell RNA-sequencing analysis and the evidence for subset of aEPDC recruited during atrial remodeling

Explants of rat atria from healthy (sham) or HF rat were cultured until epicardial cells migrated outside the explants. Cells were then sub-cultured for one passage and used for single cell RNA-sequencing.

We used a non-linear dimensionality reduction method (t-stochastic neighbor embedding t-SNE) to plot the data. In rat atria epicardium, ten clusters were defined in differentially expressed genes in aEPDCs (Fig. 6a, Supplementary Table 5). Two major populations expressing *WT1^+^* and *PDGFRβ^+^*, respectively, were clearly identifiable in both healthy and HF samples (Fig. 6b). Interestingly the *WT1^+^* population expressed *PDGFR*α and *ATGR1* (Fig. 6b). Some cells at the border of the fibroblast cluster facing the epicardial *WT1^+^* cluster highly expressed both *ATGR1* and *PDGFR*α (Fig. 6b). No significant difference in expression of *WT1*, *ATGR1*, *FSTL1* or *NPR*A is detected in epicardial cells from healthy or diseased rat hearts (Supplementary Fig. 4). 3D Tsne1 revealed a segregation of cells between sham and HF samples (Fig. 6c).

**Fig. 6.**
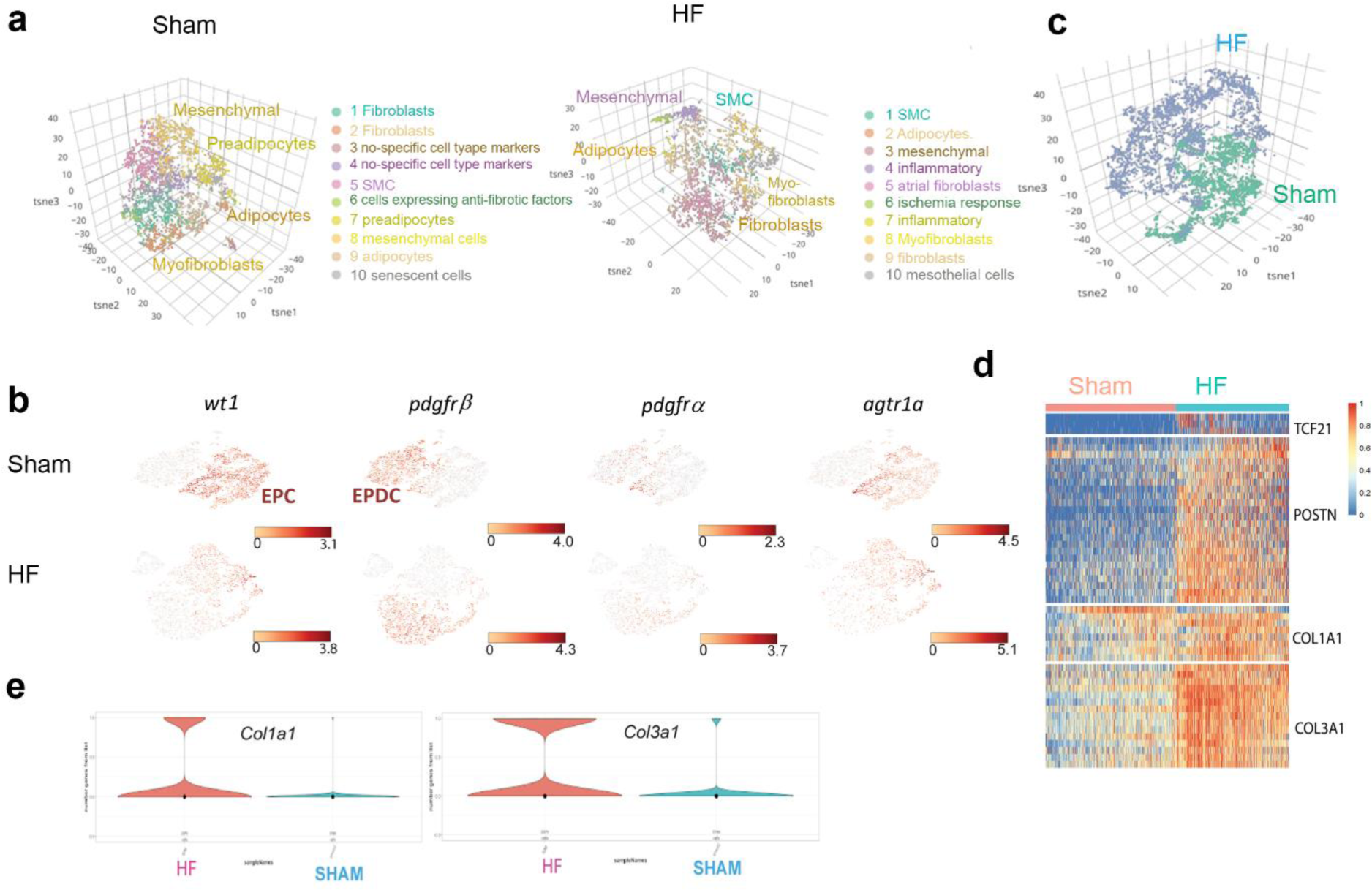
Differential transcriptional signatures aEPDCs at a single cell level between sham and diseased atria. **(a)** 3D Tsne Plots of sham and heart failure (HF) rat aEPDCs. **(b)** Tsne plots of sham or diseased (HF) epicardial (EPC) and epicardial-derived (aEPDCs) cells for the four genes (*WT1*, *PDGFR*β, *PDGFR*α, and *AGTR1*α. **(c)** 3D tsne plot of sham and HF rat aEPDCs. **(d)** Heatmap of the sham and HF aEPDCs expressing *TCF21*, *POSTN*, *COL1A1* and *COL3A1*. **(e)** Violin plots of *COL1A1* and *COL3A1* expressing cells in sham and HF aEPDCs.

On the other hand, comparison by t-SNE of the aEPDC populations in both healthy and diseased atria revealed major differences. A heatmap of five modules of differentially expressed fibrosis-relevant genes (*S100A4, COL1A1, COL3A1*) plotted from healthy and remodeled rat atria aEPDCs revealed major differences in the extent of fibrosis in each cell population (Supplementary Fig. 5). Violin plots of *COL1A1* and *-3A1,* two markers of fibroblast ECM confirmed fibrosis in HF aEPDCs (Fig. 6e, Supplementary Fig. 5).

Next, we further investigated the fibrosis process in HF aEPDCs. Clustering of HF EPDCs showed the heterogeneity of the cell population (Fig. 6a). We first applied a linear inference trajectory analysis to both sham and HF rat aEPDCs. This analysis revealed that cell transcriptomic profiles segregated the two aEPDCs populations (Supplementary Fig 6). In order to track the fibro-fatty pathological process of diseased atria, we thus performed linear trajectory inference on the HF rat aEPDCs (Fig. 7a,b). This analysis revealed that few aEPDCs at the starting point of the trajectory (cluster 10, mesothelial cells) still expressed mesothelial markers such as *MSLN.* Then, aEPDCs transitioned from adipogenic (*BMP4, LUM, UCP2*) to fibrotic (*COL1A1, COL3A1, FN1*) cell types (Fig. 7b,c, Supplementary Fig. 7, Supplementary Fig. 8). Adipogenic markers *RACK1* has been reported as an inducer of PPAR-γ and C/EBP-β, and both *RACK1* and *BMP4* are involved in adipogenesis^24, 25^ as well as the elongation factor *EEF2* reported to be a mediator of adipogenesis^26^ (module 1, Fig. 7c, Supplementary Fig 7, and Supplementary Table 5). Modules 2 and 3, included cells undergoing the pathological process of adipocyte to fibroblast transition that takes place under hypoxia in the likely presence of an inflammatory phenomenon^27^ (Fig. 7c, Supplementary Fig 7, and Supplementary Table 5). Interestingly, *CTGF,* an inhibitor of adipogenesis^28^, was increased in module 2, an index of the transition from adipose to fibrotic tissue. Finally, module 4 pointed to a feedback loop leading to a *de novo* engagement of remaining epicardial mesothelial *MSLN6+* cells through a process of EMT (*MFAP5*) toward adipogenesis (*COL6, CLMP)* and inflammatory fibrosis (*CCL2, CCL7, IL33*) (Fig. 7c, Supplementary Fig 7, and Supplementary Table 5).

**Fig. 7:**
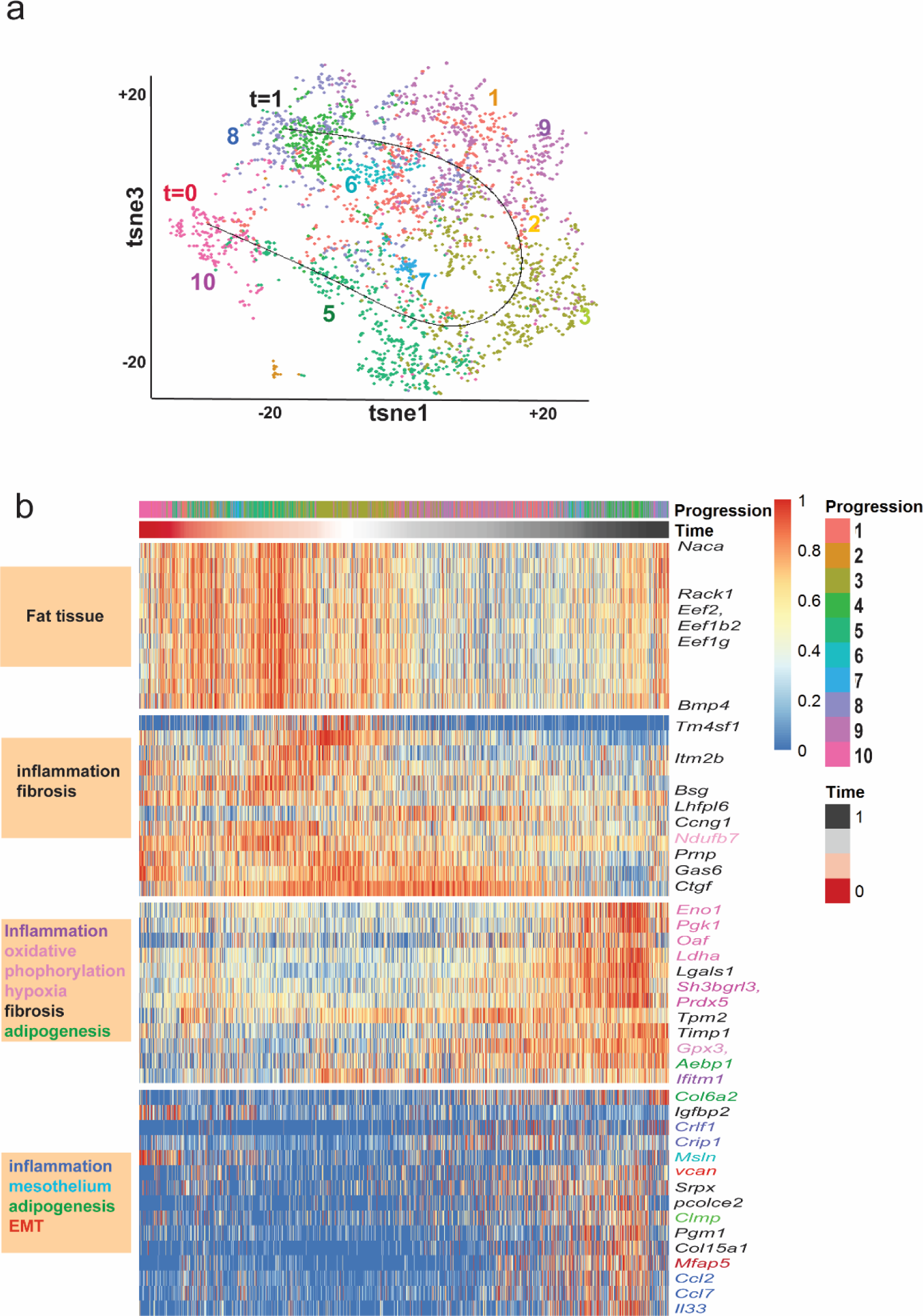
Single cell analysis of aEPDCs trajectory during atrial remodeling in rat model of heart failure (HF). **(a)** Tsne plots of clusters processed by Cell Ranger (10x Genomics). **(b)** Tsne inference trajectory of cells within the diseased aEPDC clusters (Scorpius algorithm). The pseudo-temporal ordering computational method allowed to track the fatty-fibrosis transition of cells together with an inflammatory process. **(c)** Heatmap of Inference trajectory of cells within the HF aEPDC (Scorpius algorithm). Genes are indicated on the right and colored according to the biological process indicated on the left boxes. Lines without genes (module 1) are genes without names just annoted in Ensembl (http://www.ensembl.org/index.html).

## Discussion

The accumulation of adipose and fibrosis tissue in the subepicardium has been recognized as an important determinant of the formation of the substrate of AF and understanding how the disease substrate develops is a key clinical issue. Herein, we found that following epicardial EMT, adult EPDCs maintain an adipogenic potential in epicardial layer and that they can transit to a fibrotic phenotype in response to various stimuli.

The origin of myofibroblasts and fibroblasts involved in extracellular matrix remodeling of the myocardium is still debated^22, 29^. In adult mouse ventricles, challenged by transaortic constriction or ischemia, most fibroblasts expand from developmentally-derived resident fibroblasts^22, 29–31^. Furthermore, following acute myocardial ischemia, epicardium is also activated and directly contributes to fibroblasts in the subepicardial area^2^. We provide evidence that in the atria too, epicardial expansion results in *de novo* fibroblast generation. First, cells co-expressing markers of epicardial progenitors and fibroblasts can be detected in the subepicardium of human and rat diseased atria, in agreement with a recent study^17^. Second, a number of fibroblasts proximal to the remodeled adult atrial epicardium of WT1*^CreERT^*^2^ ;ROSA26*^tdT+/-^* mice was labelled, pointing to their epicardial origin. Along these lines, it has been reported that resident epicardial precursors from the right atrial appendages can differentiate into fibroblasts, especially when obtained from AF patients^18^.

Distinct signaling pathways could operate as a switch to regulate the recruitment and fate of these epicardial progenitor cells. Indeed, several growth factor signaling pathways have been implicated in the induction of EMT, the recruitment of epicardial cells and their differentiation into mesenchymal cell lines. For instance, PDGF acts on the transcription factor Sox9 as a downstream target of PDGF stimulated EMT^23^. We reported previously that ANP is a potent adipogenic factor for aEPDCs^4^. Here, we show that during embryonic development the epicardium contains adipocyte and fibroblast precursors. Following EMT, adipocyte precursors expressing NPRA are localized in epicardium layer and AGTR1^+^ fibroblast precursors are found in sub-epicardium layer. In adult, we found that Ang-II induced differentiation of aEPDCs into fibroblasts by up-regulating its receptor, AGTR1, and by activating the canonical Smad2/3 pathway^32, 33^. Similarly, ANP up-regulated NPRA, and activated signaling resulting in adipocyte differentiation of aEPDCs^4^. Our data further suggested a positive feedback of the agonists on their signaling pathway by regulating expression of their own receptors to secure a long-lasting response of cells to the agonists. Opposite effects of Ang-II and ANP on fibrogenesis and adipogenesis, respectively, have been already reported. For instance, Ang-II inhibited differentiation of human pre-adipocyte cells^34^. An anti-fibrogenic effect of ANP has been described and attributed to the inhibition by the peptide of myofibroblast differentiation and fibroblasts activation^35^. Finally, Ang-II down regulates NPR1 expression at the transcription level^36^. These data suggest that from the embryo the epicardium progenitors could be pre-defined into different fates and their re-activation is depend of local stimuli.

Using scRNA-seq, we were able to provide novel insight into aEPDC heterogeneity and the relationships between aEPDC populations. Single cell RNA-seq analysis revealed six clusters of atrial epicardial cells and aEPDCs, including two predominant epicardial *WT1^+^* and *PDGFRβ^+^* clusters, and identified adipogenic and fibrogenic potential of these cells. Another major result provided by the scRNA-seq analysis of aEPDCs from diseased atria, is the process of a transition of epicardial cells from adipose and then fibrotic tissue. Our scRNA-seq data confirmed that a *de novo* EMT of remaining more resistant epicardial cells in the culture could be triggered by hypoxia and inflammatory processes (module 4, Fig 7c), giving rise to adipose tissue and further fibrosis as reported during obesity^37^. Indeed, linear trajectory inference suggested a progression of the disease from transient accumulation of fat tissue towards fibrosis. However, our data do not exclude that EPC undergoing EMT directly differentiate into fibroblasts.

This self-help mechanism originating from the epicardium and resulting in the generation of fibroblasts could explain why thin epicardial layer co-exists with thick and fibrotic epicardial area. This histological observation suggests that local mechanisms regulate the activation the epicardium resulting in focal accumulation of adipocytes and fibroblasts. An additional argument for local activation of the epicardium is that aEPDC-derived fibroblasts are observed only in the atria but not in the ventricle of WT1*^CreERT2+/-^*;Rosa26*^tdT+/-^* mice. Moreover, the thick and fibrotic remodeled epicardium predominates in atria from aged patients during clinical setting associated with atrial dilatation suggest that activation of the epicardium is a low grade process that contributes with time to the progression of atrial remodeling and the formation of AF substrate. Taken together, these histological and biological findings point to the presence of a subpopulation of aEPDCs that are pre-engaged in adipogenic or fibrogenic lineages, in the subepicardium and ready to be recruited in response to various stimuli, resulting, with time, in different degrees of fibro fatty infiltrates.

## Conclusion

We found that subsets of cells derived from epicardial progenitors and confined into the subepicardium regulate, the balance between AT expansion and fibrosis accumulation in the atrial myocardium. Multiple factors coupled to signaling pathways can contribute to the recruitment of these “post EMT EPC” from their subepicardial “niche”. Our study provides a biological basis for the slow and low noise remodeling of the atrial subepicardium which progresses with time or during chronic cardiac diseases. It remains to determine when this apparent aging phenomenon becomes pathological and can contribute to the substrate of AF.

## Acknowledgement

We are grateful to Orestis Faklaris from Jacques Monod Institute for technical support with polarized light and Florence Deknuydt from cardiometabolism and nutririon for technical support with fluorescence-activated cell sorting and flow cytometer.

## Affiliations

From Sorbonne universités, Faculté de médecine UPMC, Paris, France (N.S., N.M., D.A.T., G.D., M.B., P.L., S.N.H.); Aix Marseille Univ, INSERM, MGG, U 1251, Marseille, France (T.M.M., M.P.); Department of Cardiology, Assistance Publique - Hôpitaux de Paris, Pitié-Salpêtrière Hospital, Paris, France (P.L., S.N.H.); Medical Biochemistry Institute and molecular biology, University Medical Center, Greifswald Germany (C.W., U.L.); Institute of Cardiometabolism and Nutrition, ICAN, Paris, France (N.S., N.M., D.A.T., G.D., M.B., P.L., S.N.H.).

## Contributions

N.S. contributed to the design of the experiments, conducted the experiments, analyzed data and was involved in writing of the manuscript. T.M-M. contributed to WT1Cre*^ERT2+/-^*;ROSA26*^tdT+/-^* transgenic mice, to the design of the experiments, conducted the experiments, analyzed data and was involved in writing of the manuscript. N.M. contributed to realize and characterize experimental models and analyze data. G.D. and M.B. contributed to perform histological research. B.J. contributed to analyze scRNA-seq. J.P. contributed to perform research. P.L.P. contributed to perform research and provide human samples. D.A.T contributed to analyze data. M.P. contributed to the design of the experiments, conducted the experiments, analyzed data and was involved in writing of the manuscript. S.N.H contributed to the design of the experiments and supervised and wrote of the manuscript.

---

**Supplementary Fig. 1.**
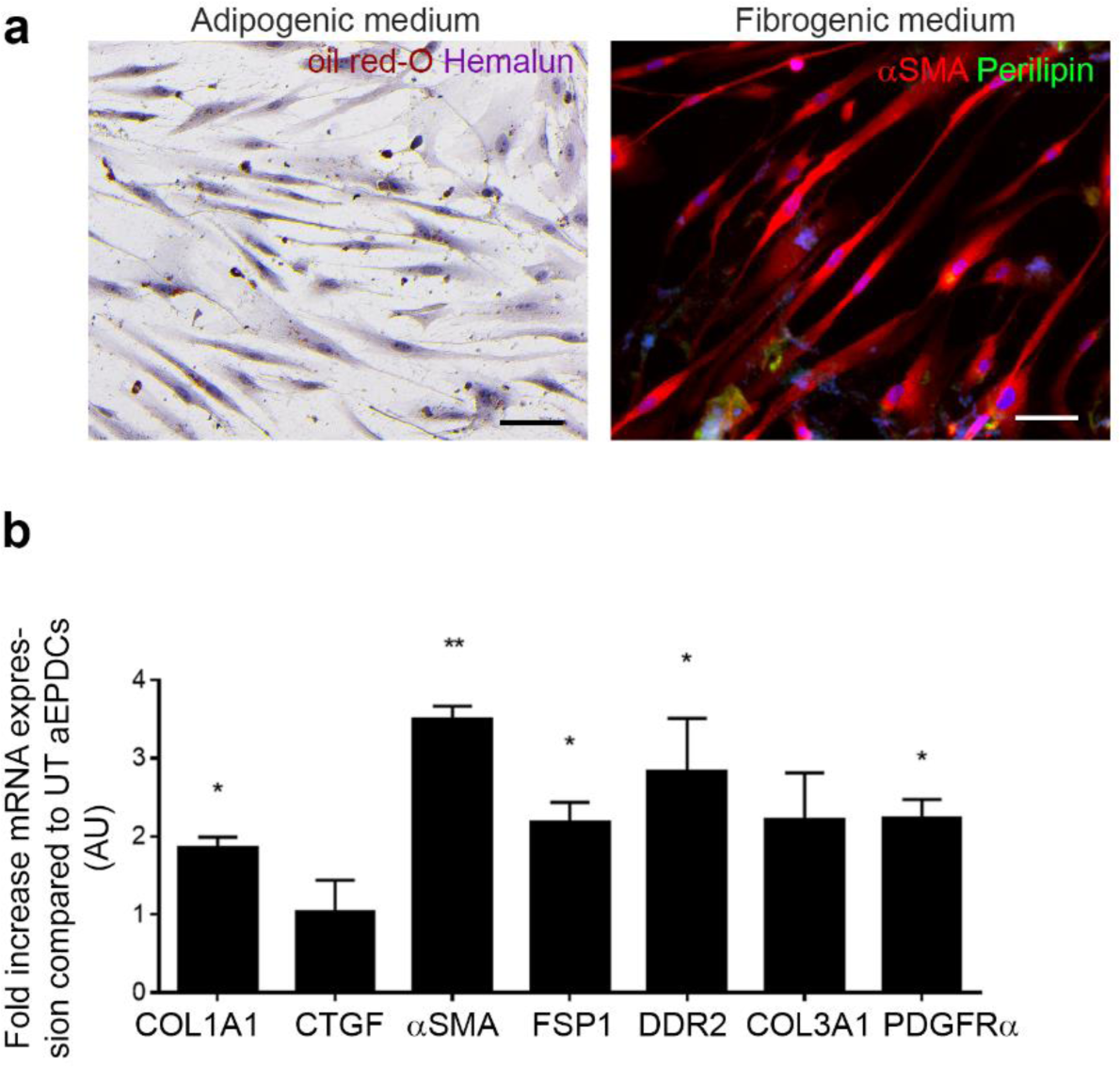
TGF-b induces fibrogenic differentiation of aEPDCs. (**a**) Representative images of aEPDCs treated with TGF-β (10 nM) for 7 days and stained with oil-red-O and counterstained with hemalun or immuno-labeled for perilipin or αSMA (n=5). Scale bar, 100 µm. (**b**) Expression of fibrosis genes in aEPDCs treated with TGF-β (10 nM) for 7 days compared to untreated (UT) control. Data are expressed as mean ± SEM of n=4 independent experiments. *, P < 0.05, **, P <0.01, one-way ANOVA with Bonferroni’s post hoc test.

**Supplementary Fig. 2.**
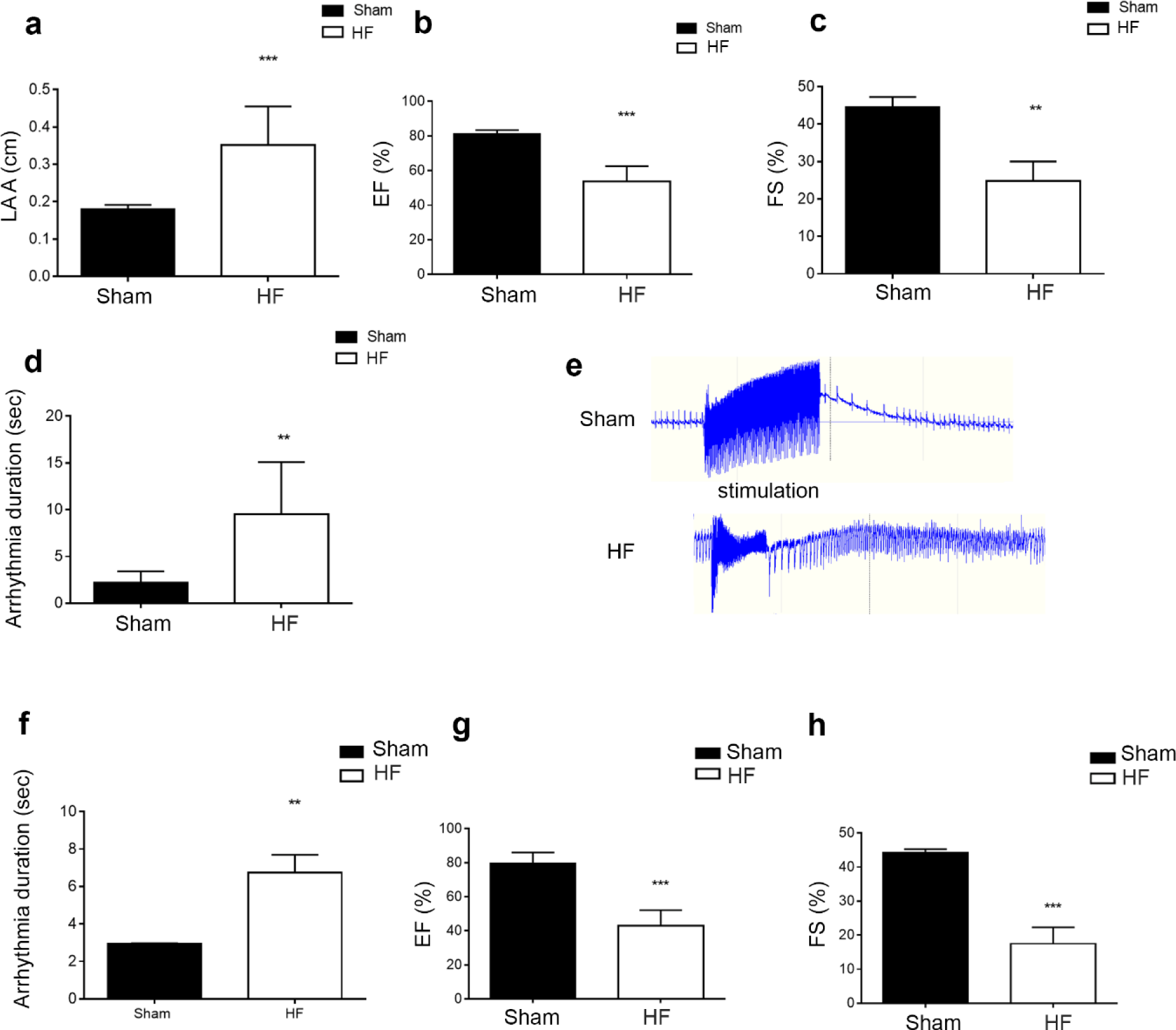
Vulnerability of AF in rat or mouse model of atrial remodeling associated with HF. In rat 2-months post-MI (n=34) or sham (n=12) (**a**-**e**) or in mouse 2-months post-MI (n=6) or sham (n=10) (**f**-**h**), histograms represent left atria area (LAA) (**a**), ejection fraction (EF) (**b**,**g**), fractional shortening (FS) (**c**,**h**) or arrhythmia duration (**d**,**f**). (**e**) Paroxysmal atrial fibrillation episodes in rats 2-months post-HF. Each data are expressed as the mean ± SEM of independent experiments. **, P <0.01, *** P <0.001, unpaired Student’s t test.

**Supplementary Fig. 3.**
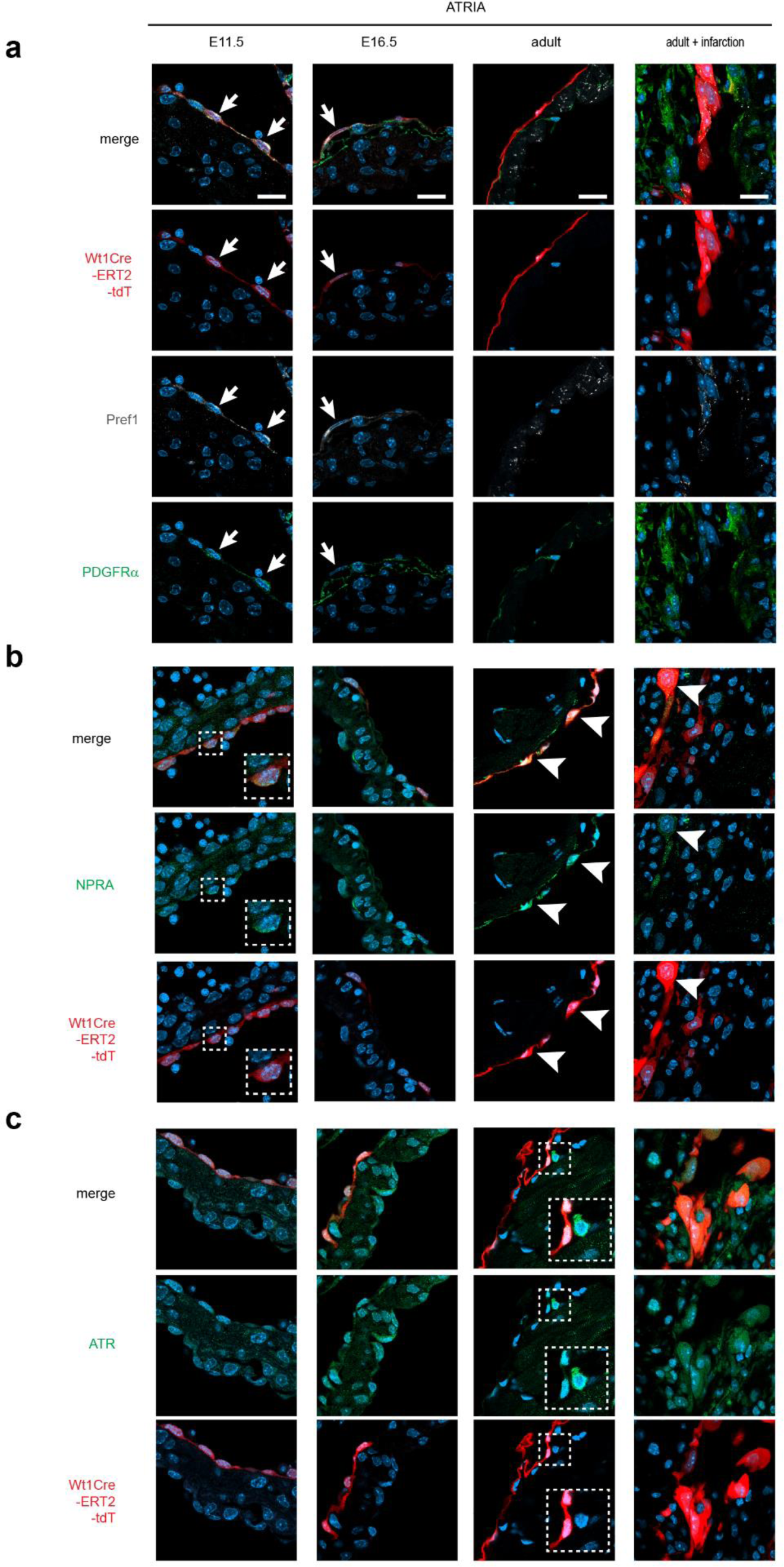
Heterogeneity of atria epicardium in mouse model of atrial remodeling associated with HF. Representative images of heterogeneity of Pref1 and PDGFRα (**a**), NPRA (**b**) or ATR1 (**c**) expression in E11.5, E16.5, adult sham or HF atrial epicardium (**a**), Scale Bar, 10µm, 50 µm.

**Supplementary Fig. 4:**
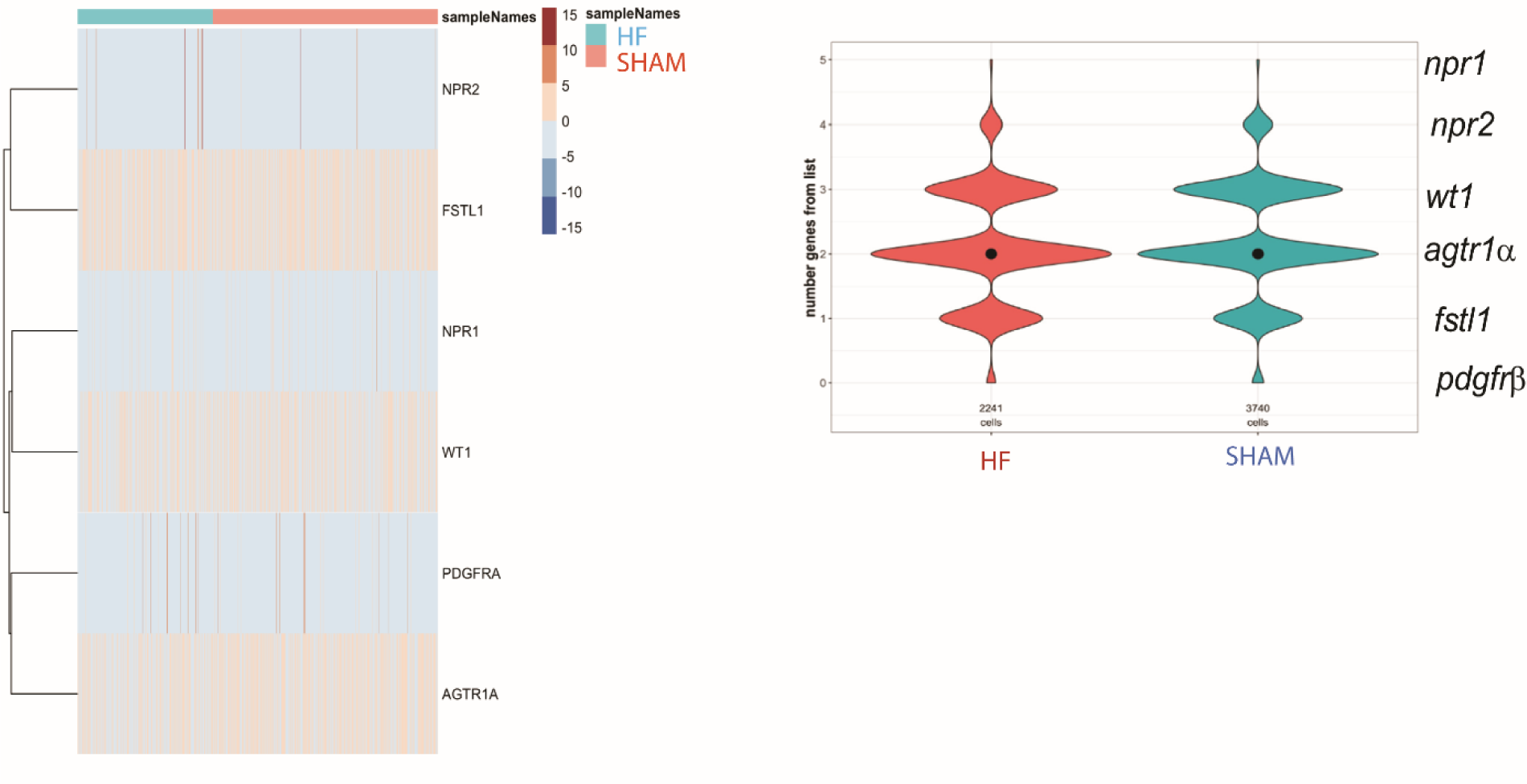
gene profile of rat epicardial cells from healthy and diseased (HF) rat atria. Left panel: Heat map of epicardium specific genes in both sham and HF dataset. Right panel: co-expression violin plot of 5 genes in both sham and HF dataset.

**Supplementary Fig. 5:**
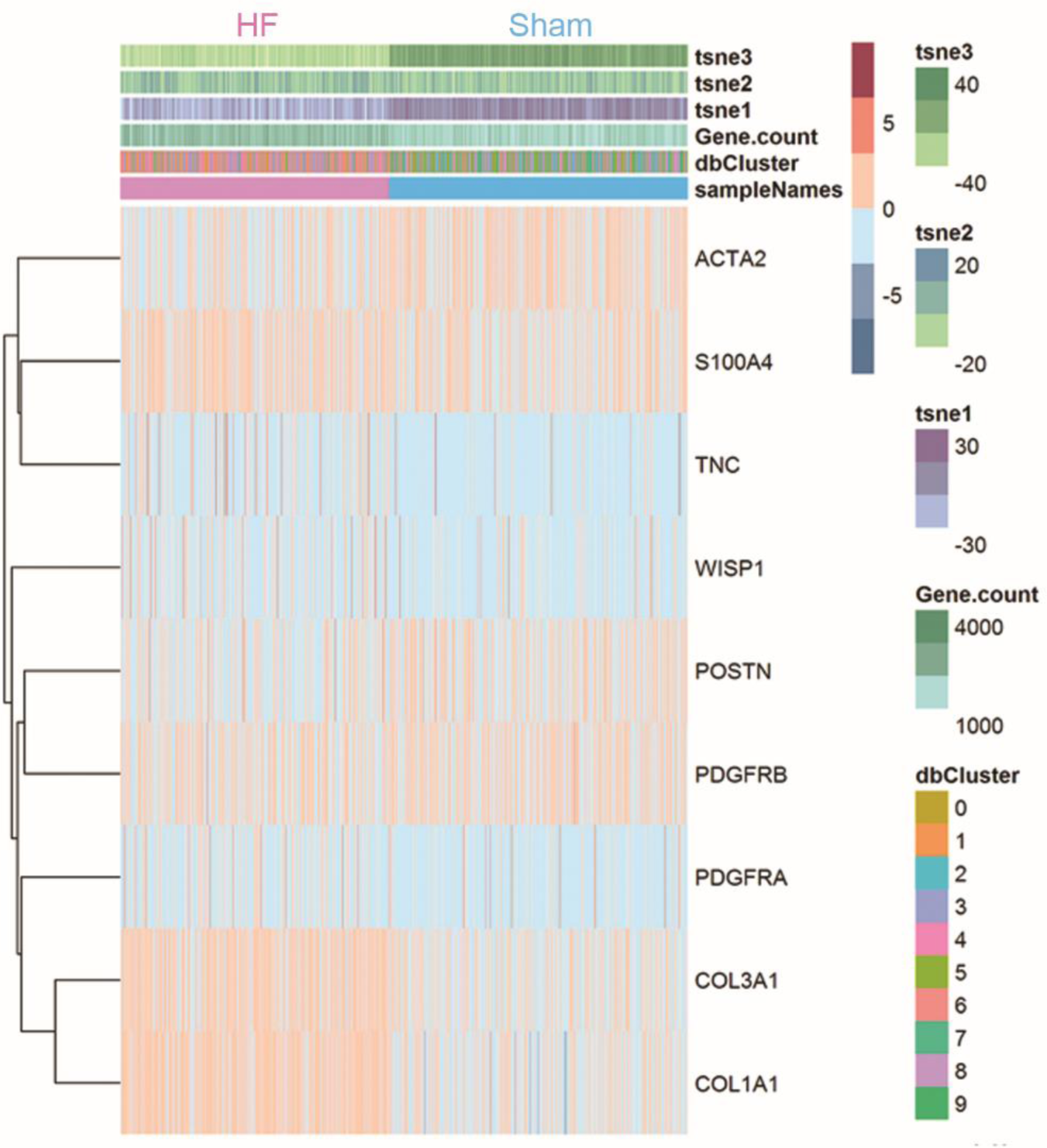
Heat map of fibrotic genes in healthy (Sham) and diseased (HF) rat atria aEPDCs.

**Supplementary Fig. 6:**
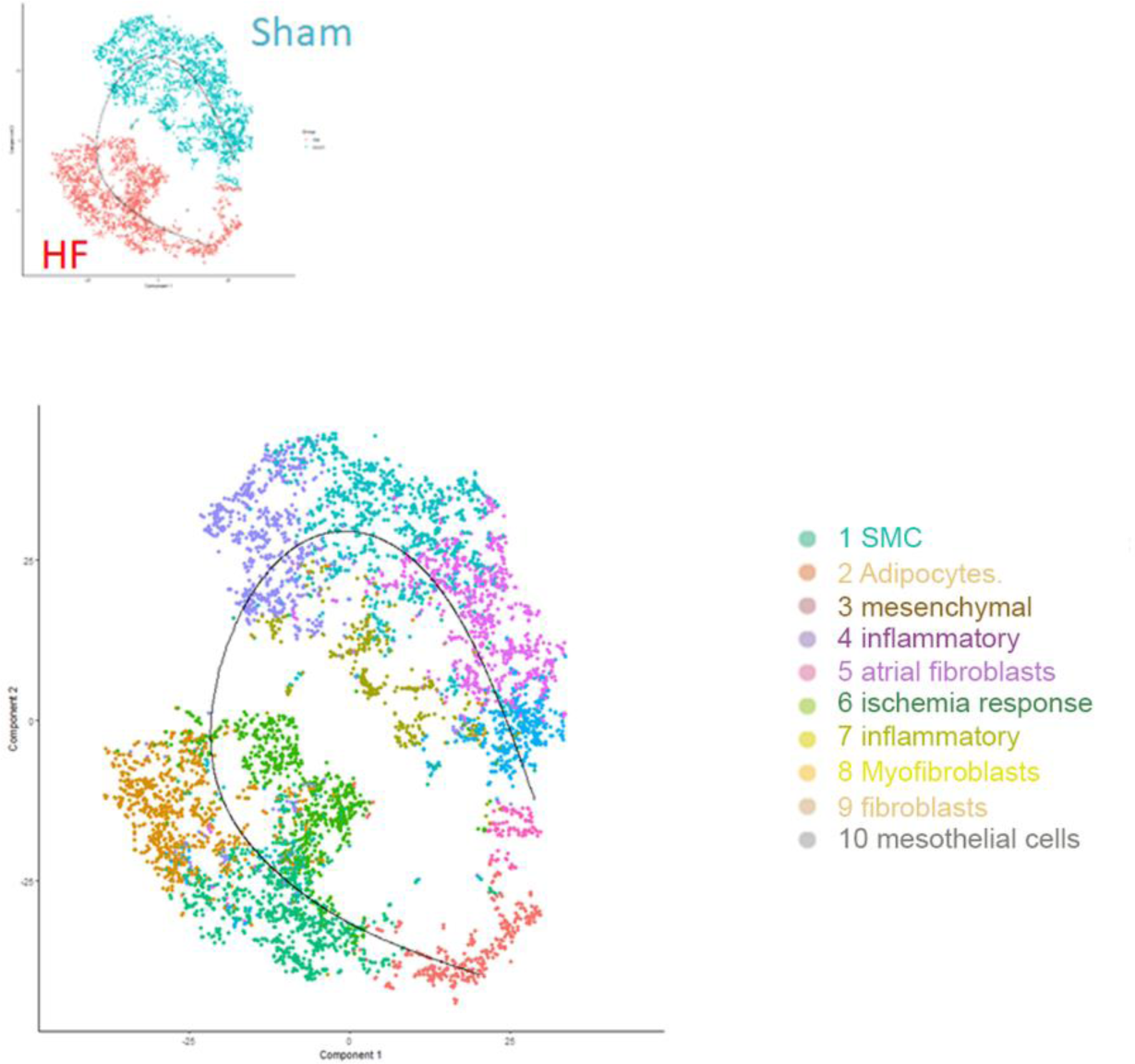
Trajectory from healthy (Sham) to diseased (HF) rat atria aEPDCs.

**Supplementary Fig. 7:**
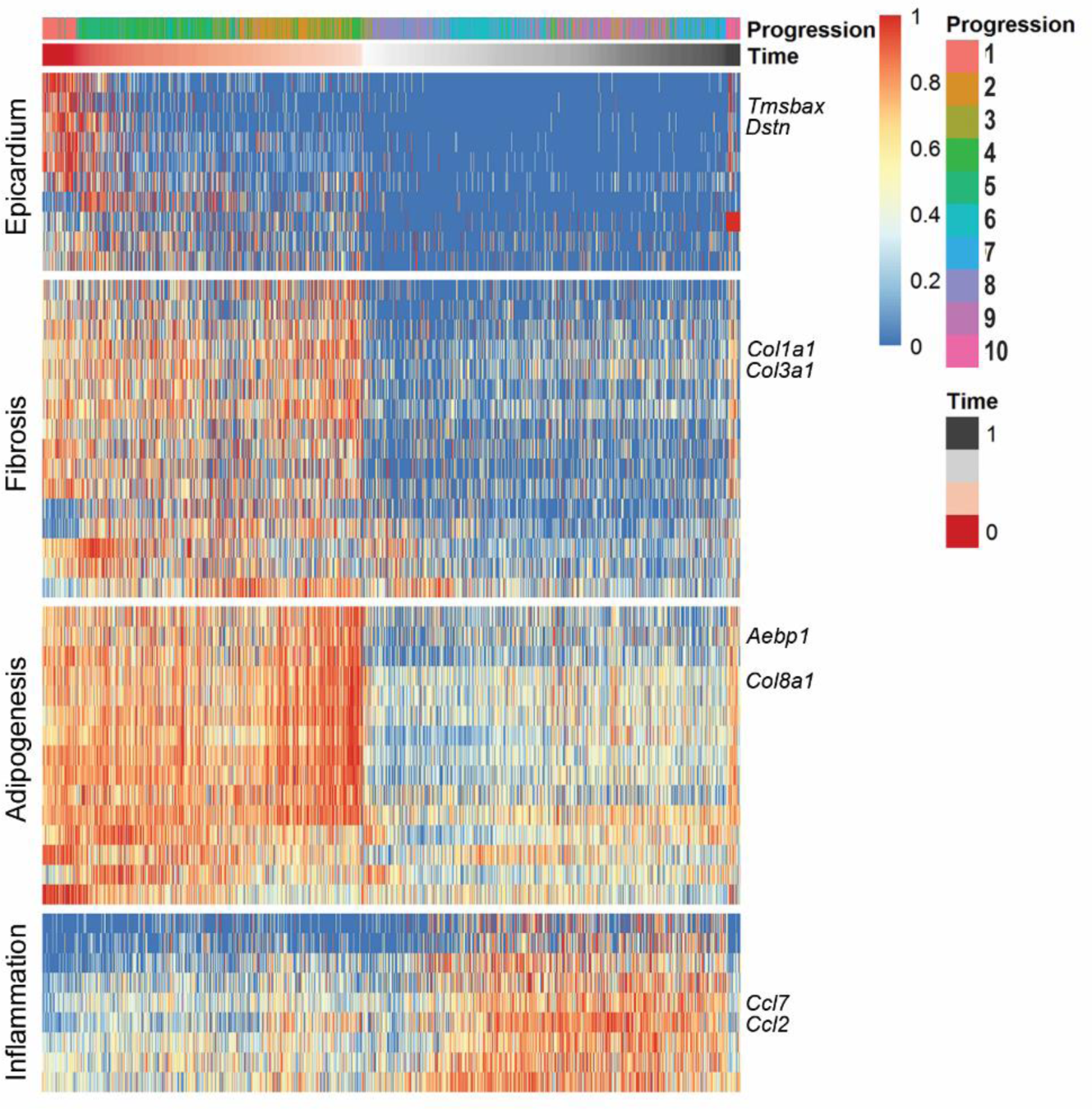
Trajectory influence heat map of healthy (Sham) and diseased (HF) rat atria aEPDC. Data are analyzed wit Scorpius logarithm.

**Supplementary Fig. 8.**
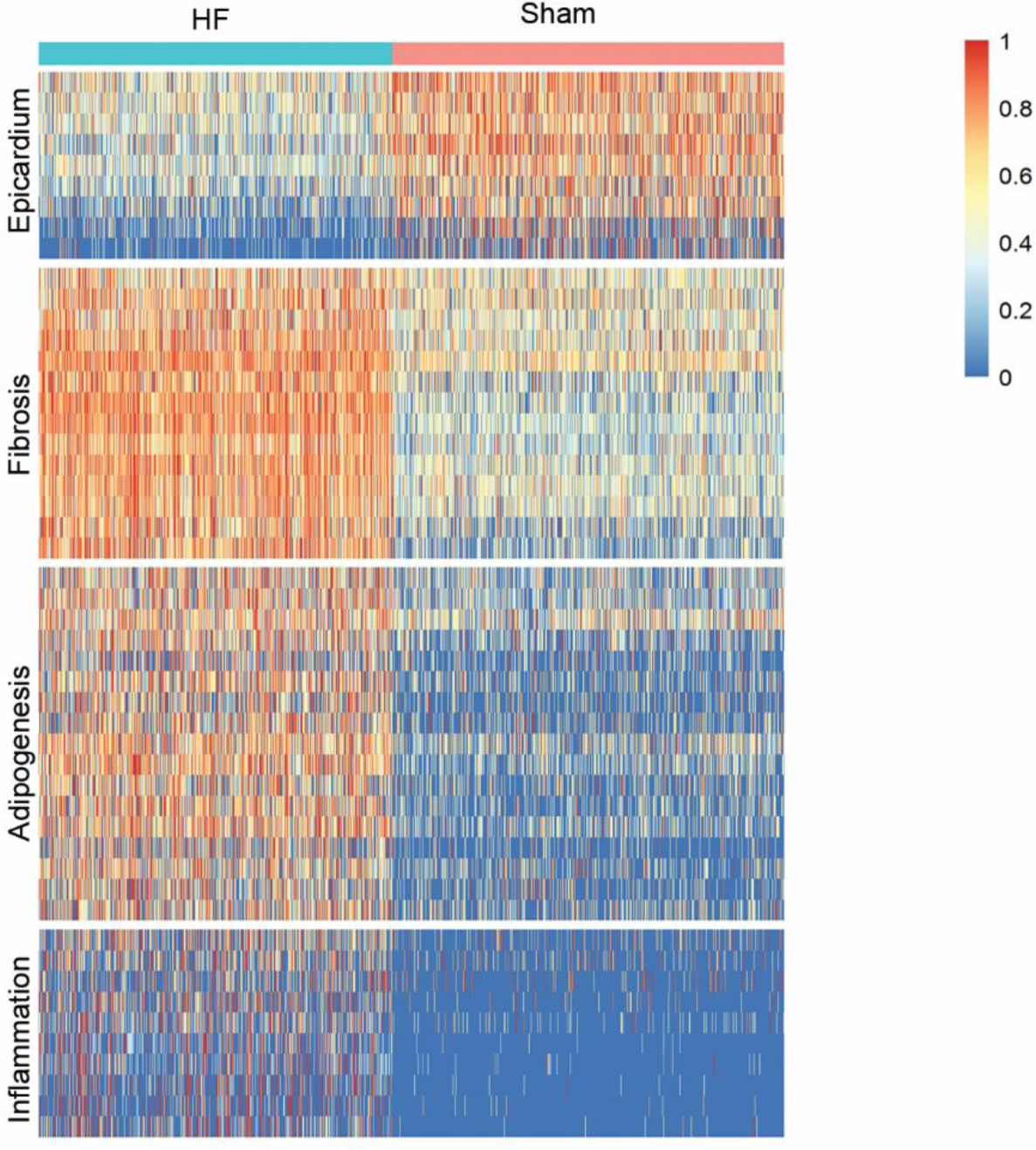
Heat map of adipogenic and fibrotic genes expression in both healthy and diseased (HF) rat atria aEPDCs. Data are analysed wit Scorpius logarithm.

**Supplementary Table 1.** Clinical parameters of patients. BMI, body mass index; LVEF, left ventricular ejection fraction; AF, Atrial fibrillation; VMR, Mitral valve replacement.

| Clinical outcomes for | Histology | EPDCs | Secretome |
| --- | --- | --- | --- |
| Number of subjects, n | 109 | 15 | 10 |
| Age, y (mean ± SD) | 69 ± 8.5 | 67 ± 9 | 66 ± 6 |
| Sex, M/F | 64/45 | 8/7 | 2/8 |
| BMI, kg/m<sup>2</sup> (mean ± SD) | 26 ± 3 | 25.5 ± 4.2 | 27.3 ± 1.2 |
| AF | 31 | 4 | 3 |
| LEVF <50%, n | 84 | 2 | 2 |
| Coronary bypass, n | 41 | 7 | 6 |
| VMR, n | 20 | 8 | 4 |

**Supplement Table 2.**
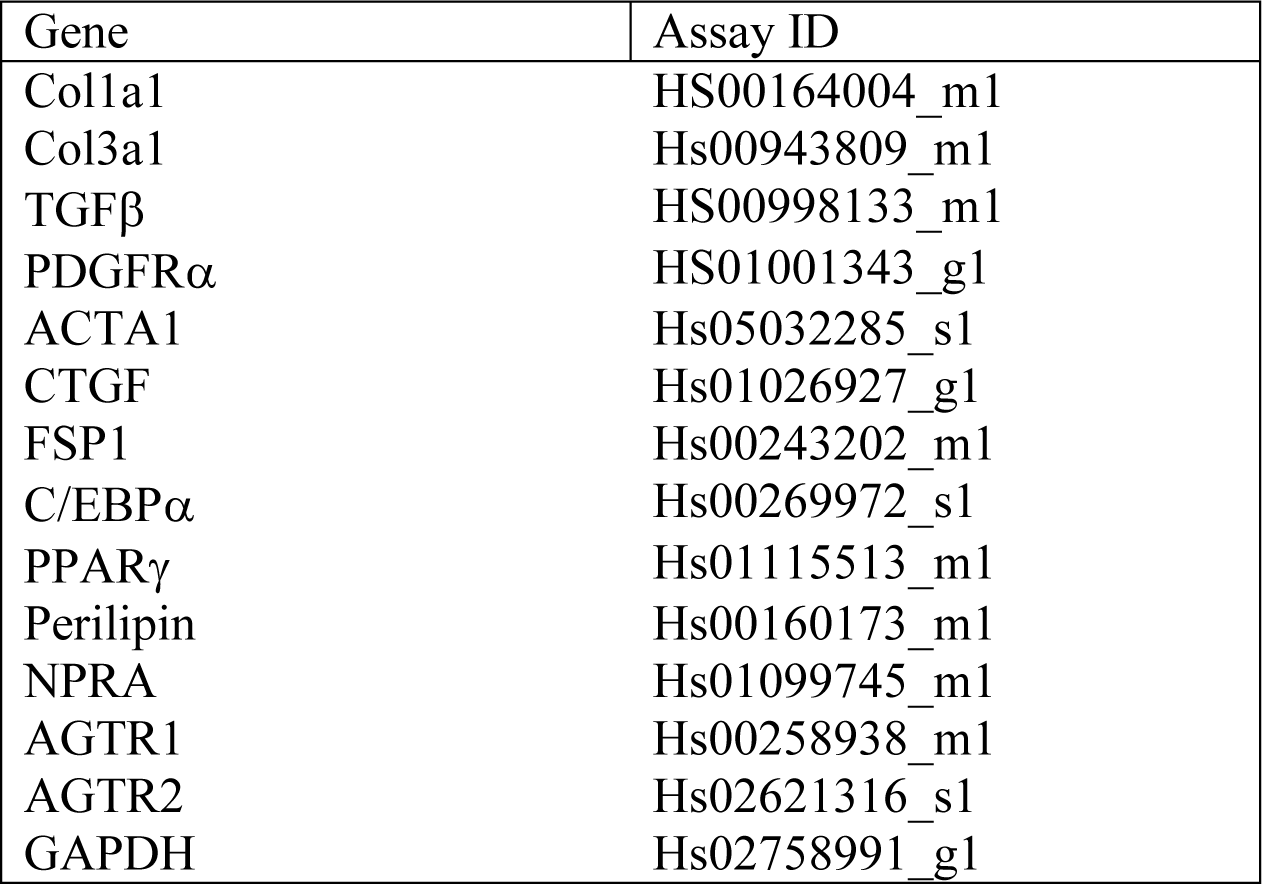
Taqman Human gene probes.

**Supplementary Table 3.** (**a**) Cell number per field in human atrial tissue. (**b**) Cell number per field in rat atrial tissue.

| Cells, (n) | Mean ± SEM | Percent (%) | *P value* |
| --- | --- | --- | --- |
| Dapi | 306±120.9 | 100 | - |
| WT1(+) | 46±19.5 | 15±2 | P<0.0001 |
| PDGFRα(+) | 44±12 | 10.3±2.1 | P<0.0001 |
| αSMA(+) | 15±4.5 | 8.4±1.1 | P<0.0001 |
| PDGFRα(+)WT1(+) | 25±11 | 8.4±0.4 | P=0.0579 |
| αSMA(+)<br>WT1(+) | 9±5 | 3.1±0.1 | P=0.103 |

(b) Cell number per field in rat atrial tissue
| Cells, (n) Mean ± SEM | MI | Sham | *P value* |
| --- | --- | --- | --- |
| Dapi | 804±237 | 580±205 | P<0.05 |
| WT1(+) | 80±26 | 31±9 | P<0.05 |
| PDGFRα(+) | 100±29 | 26±8 | P<0.05 |
| TCF21(+) | 150±40 | 74±31 | P=0.0714 |
| Pref-1(+) | 11±2 | 17±6 | P=0.2460 |
| PDGFRα(+)TCF21(+) | 55±17 | 13±5 | P<0.05 |
| Pref-1(+)<br>WT1(+) | 11±2 | 15±6 | P=0.5317 |

**Supplementary Table 4.** Association of biological parameters with clinical outcomes. LVEF, left ventricular ejection fraction; AF, Atrial fibrillation; VMR, Mitral valve replacement; (ye), years.

|  | Adipose tissue/<br>Epicardial Fibrosis |  |  |
| --- | --- | --- | --- |
|  | LVEF<50 | LVEF>50 | P value |
| age (ye) |  |  |  |
| < 70 | 0.52 | 0.55 | 0.8289 |
| ≥ 70 | 2.53 | 0.59 | 0.0389 |
|  | VMR<sup>-</sup> | VMR<sup>+</sup> | P value |
| age (ye) |  |  |  |
| < 70 | 0.48 | 0.75 | 0.595 |
| ≥ 70 | 0.84 | 0.33 | 0.0425 |
|  | Bypass<sup>-</sup> | Bypass<sup>+</sup> | P value |
| age (ye) |  |  |  |
| < 70 | 0.96 | 0.47 | 0.323 |
| ≥ 70 | 0.44 | 0.96 | 0.219 |
|  | AF<sup>-</sup> | AF<sup>+</sup> | P value |
| age (ye) |  |  |  |
| < 70 | 0.59 | 0.19 | 0.764 |
| ≥ 70 | 0.86 | 0.32 | 0.2119 |

**Supplementary Table 5.** Dataset from scRNAseq analysis.

